# C26 and CT26 colorectal cancer models exhibit divergent cachexia phenotypes, intramuscular inflammation, and protein turnover signaling

**DOI:** 10.64898/2026.04.21.719997

**Authors:** Xinyue Lu, Hanan Tlais, Hamood Rehman, Charles P. Olson, Abigail L. Hartz, Nina R. O’Tain, Ashtyn N. Martens, Tingting Ju, Vandré C. Figueiredo, James F. Markworth

**Author notes:** **Corresponding author:** James F. Markworth, Ph. D.

## Abstract

Colorectal cancer (CRC) cachexia induces skeletal muscle dysfunction, impeding quality of life and worsening cancer prognosis. Multiple preclinical models, including the widely used mouse model of subcutaneous inoculation with the C26 colorectal carcinoma cell line, have been developed to study the biological mechanisms of CRC cachexia and elucidate potential new treatments. It has been proposed that a distinct cell line of the same origin, namely CT26, is relatively non-cachectic. However, studies evaluating the relative potential of C26 and CT26 cells to induce cancer cachexia in parallel have been limited. The differences in the biological mechanisms by which C26 and CT26 impact skeletal muscle mass and function have also not been fully elucidated. In the current study, we investigated the differential capacity of C26 and CT26 to induce cancer cachexia using both an *in vitro* cancer-muscle cell co-culture and an *in vivo* syngeneic mouse model. Our results show that both C26 and CT26 cells induced significant atrophy of murine C2C12 skeletal myotubes. In the mouse model, while C26 and CT26 both reduced skeletal muscle mass and fat mass, only C26 tumors led to loss of body weight and impaired skeletal muscle force output. We further show that C26 tumors express a higher level of interleukin-6 and C26 tumor-bearing mice exhibit greater muscle inflammation than CT26 tumor-bearing mice. In addition, mice bearing C26 and CT26 tumors showed differential regulation of the innate immune responses and muscle protein turnover. Overall, our data suggests that although both C26 and CT26 cells do exhibit cachectic effects, C26 cells induce greater loss in body weight, fat mass, skeletal muscle mass, and physical function via promoting chronic inflammation and deregulating protein balance of skeletal muscle.

## INTRODUCTION

Cancer cachexia is a multifactorial syndrome characterized by progressive loss of body weight, skeletal muscle mass, adipose tissue, and physical function (1). As the second leading cause of cancer-related mortality, colorectal cancer (CRC) is projected to impact over 150,000 members of the US population in 2026, with a growing burden of incidence in adults under 50 years old (2). Over 30% of CRC patients develop cancer cachexia, which is closely associated with cancer morbidity and mortality (3–5). During CRC progression, the loss of skeletal muscle mass is an independent predictor for worsened responses to anti-cancer treatments and poorer CRC prognosis (6, 7). However, despite active research on anti-cachexia interventions, medical strategies to alleviate cancer cachexia in CRC patients are currently limited (8).

Multiple preclinical models have been developed to investigate the biological mechanisms of CRC cachexia and discover potential novel anti-cachectic interventions. One of the most prevalent pre-clinical models of cachexia is that induced by colon-26 (C-26) colorectal carcinoma cells (9, 10). First established by Corbett et al. 1975, the C26 cell line was originally derived from female BALB/c mice receiving the colon carcinogen N-nitroso-N-methylurethane (NMU) (11). C26 tumors were identified as an undifferentiated colon carcinoma that was highly tumorigenic (11, 12). Extensive subsequent research has shown that tumor-derived factors secreted from C26 cells result in profound muscle wasting in the terminally differentiated cultures of the C2C12 murine skeletal muscle cell line (13, 14). C26 cells have also been widely adopted in animal models, with various studies testing subcutaneous, intravenous, or intramuscular injections induce tumor growth and cancer cachexia in both BALB/c and CD2F1 mice (10, 15–20). Early tumorigenicity research showed that injection of a dose of 1 × 10^5^ C26 cells led to a 100% tumor induction by two weeks post-inoculation (12). Mice bearing C26 tumors display decreased body weight, skeletal muscle mass, and adipose tissue mass, as well as hypoglycemia and hepatic dysfunction (10, 21). Mechanistically, cachexia in C26 tumor-bearing mice has been associated with increased muscle inflammation, impaired anabolism/increased catabolism, disturbed membrane integrity, and impaired the mitochondria function (18, 22–25).

Another CRC cell line of the same origin, termed CT26, has been relatively less studied. The CT26 cell line was also originally derived from NMU-induced colon tumor in BALB/c mice, yet there is no record showing when and how the CT26 cell line diverged from C26 cells, and some studies even use the terms C26 and CT26 interchangeably (26). Recent research on tumor immunogenicity of mice bearing C26 versus CT26 carcinomas demonstrates distinct differences in genetic mutations and tumor immune infiltration of the two cell lines. While both cell lines have KrasG12D mutation, genomic profile demonstrates that CT26 cells possess unique mutations in oncogenes (e.g., *Birc5* and *Mki67*) and tumor suppressors (e.g., *Cdkn2a*) not observed in C26 cells (27, 28). On the other hand, C26 cells have a higher level of Wnt signaling activity than CT26 cells (29). Furthermore, *in vivo* studies have demonstrated that while CT26 tumor tissues are infiltrated with more cytotoxic T cells, dendritic cells, and natural killer (NK) cells, that C26 tumors rather exhibit lower T cell infiltration and higher resistance to anti-PD1 therapy (29).

Several studies have reported body weight loss and muscle wasting in CT26 tumor-bearing mice (30–33). Nevertheless, other groups have proposed that CT26 cells might be relatively incapable of inducing cancer cachexia (34–36). The inconsistent observations of investigators employing the CT26 model could perhaps be attributed to the differences in sources of CT26 and the physiological characteristics of the hosts. In addition, tumor progression and the efficacy of inducing cachexia could potentially be impacted by the site of tumor inoculation (37, 38). However, research directly comparing C26 and CT26 cells in parallel for their potential of inducing cancer cachexia and possible differences in biological mechanisms has been limited to date (34, 36).

It is worth mentioning that most prior C26 or CT26 inoculation studies were done in young mice of 6-8 weeks (39, 40). At this young age, mice are still typically growing and thus gaining body weight and increasing skeletal muscle fiber size (41, 42). Although prior research has shown that C26 successfully induces cancer cachexia in both young developing mice (8 weeks) and middle aged adult mice (12 months), differential impacts on fiber size and proteolysis pathways was observed between age groups (43). Notably, the average onset age of CRC in human populations is over 50 years old, with an increase in early-onset CRC (35-49 years) (44). Therefore, cachexia models using young mice of 6-8 weeks might not fully represent the physiological characteristics in adults developing CRC cachexia. Thus, it might be meaningful to investigate how CRC cachexia influences skeletal muscles in older adult mice that better represent the average onset of colorectal cancer in the human population (45, 46).

In the current study, we aimed to investigate the potential differential impacts of C26 and CT26 colon carcinoma cells on skeletal muscle physiology as well as the underlying biological mechanisms. To examine possible difference between *in vitro* and *in vivo* models of cancer cachexia, we used both a cancer-muscle cell co-culture model and a mouse model of subcutaneous tumor inoculation. While both C26 and CT26 cells induced comparable atrophy in C2C12 myotubes *in vitro*, C26 cell inoculation resulted in relatively greater decrements in skeletal muscle mass and strength in BALB/c mice *in vivo*. In addition, we observed marked local overexpression of interleukin-6 (IL-6) in C26 tumor tissue, whereas the CT26 group exhibited higher immune cell infiltration in the tumor microenvironment. Immunohistochemistry and molecular analysis revealed that C26 tumor-bearing mice exhibited greater skeletal muscle inflammation associated with an overexpression of the pro-inflammatory enzyme 5-lipoxygenase (5-LOX) and a marked depletion of the anti-inflammatory/pro-resolving enzyme 12/15-lipoxygenase (12/15-LOX). C26 and CT26 tumors also showed differential regulation of the muscle protein synthesis and degradation pathways.

## METHODOLOGY

### Cell culture

The murine skeletal muscle cell line C2C12 (ATCC, CRL-1772) and two colorectal carcinoma cell lines including C26 (a gift from Dr. Kee-Hong Kim at Purdue University) and CT26 (ATCC, CRL-2638) were cultured separately in growth media consisting of Dulbecco’s modified Eagle medium (DMEM, Gibco, 11995-065) containing 10% fetal bovine serum (FBS, Corning, 35-010-CV) and antibiotics (penicillin 100 U/mL, streptomycin 100 μg/mL, Gibco, 15140-122) at 37 °C in a cell incubator with 5% CO_2_.

### Co-culture model

When reaching a confluency of 70%, C2C12 cells were passaged and seeded into 24-well co-culture base plates (Corning, 353504) in growth media at a cell density of 2.5 × 10^4^/cm^2^. Four days after plating, the C2C12 myoblasts in the base plate were induced to form myotubes by replacing the growth media with differentiation media consisting of DMEM containing 2% horse serum (HS, Gibco 26050088) and antibiotics. On the day of induction of myogenic differentiation, C26 and CT26 cells were seeded in growth media at a low density (4.17 × 10^4^/cm^2^), a moderate density (8.3 × 10^4^/cm^2^), or a high density (1.67 × 10^5^/cm^2^) into the upper compartments of co-culture plate inserts (Corning, 353095) that were placed into base plates lacking cells. Both C26 and CT26 cells were used at a low-to-moderate passage number (C26 at P13 and CT26 at P15). The two cell lines were always handled side by side and used within 7 passages after thawing from a frozen stock. C2C12 myoblasts (8.3 × 10^4^/cm^2^) were plated in the inserts as a non-cancer cell control for the co-culture experiment. Following 72 hours of myogenic differentiation of the C2C12 myotubes in the base plates, the upper co-culture inserts containing C2C12 myoblasts, C26 cells, or CT26 cells were moved to the lower compartments of the base plates. Each of the cell types in the upper compartments was 100% confluent when moved to the C2C12 myotubes. Fresh differentiation media was added to both the C2C12 myotubes in the base plates and the various cell types in the upper co-culture inserts when the co-cultures were established. C2C12 myotubes were co-cultured with C2C12 myoblast control, C26 cells, or CT26 cells for 72 hours before fixation and immunocytochemistry as described below.

### Mice and tumor inoculation

10-month-old female BALB/c mice were obtained from Charles River Laboratories and housed under specific pathogen-free conditions with ad libitum access to food and water. After one week of acclimatization, the mice were randomized to three groups with balanced starting body weights. Prior to tumor inoculation, C26 and CT26 cells were harvested side by side and resuspended in sterile saline. Both C26 and CT26 cells were inoculated at low-to-moderate passage numbers (C26 at passage 14 and CT26 at passage 12), and both cell lines were cultured for the same number of passages after thawing from liquid nitrogen (<6 passages) before use. Two groups of mice were inoculated with either C26 cells (5 × 10^5^ cells/100 μL saline per mouse, *n* = 7) or CT26 cells (5 × 10^5^ cells/100 μL saline per mouse, *n* = 7) subcutaneously in their upper right flank. Age-and sex-matched healthy control mice (*n* = 6) were injected with sterile saline alone (100 μL/mouse). Mouse condition and body weight were monitored at least three times a week, and tumor dimensions (length and width) were measured by a caliper every other day starting day 5 post inoculation. Mouse food intake was measured every two days. The animal study was approved by the Purdue University Institutional Animal Care and Use Committee (IACUC protocol #2205002271. Mice were euthanized as a humane endpoint following the criteria of IACUC policy when (1) any of the existing tumors reached over a diameter of 2.0 cm, (2) diameters of multiple tumors added up to over 3.0 cm, (3) the body condition score (BCS) reached <2/5, or (4) tumors interfered with normal functions (eating, ambulating, eliminating). Euthanasia as humane endpoints were accounted for when calculating the survival curve. Animals were euthanized on day 24 post inoculation or upon reaching humane endpoint criteria, whichever occurred first.

A second cohort of 10-month-old female BALB/c mice were also injected with saline (*n* = 5), CT26 cells (*n* = 5), or C26 cells (*n* = 5) with the same procedure described above. Mouse body weight and tumor dimensions were monitored three times a week, and food intake was recorded every two days. This additional cohort was collected 13 days post tumor inoculation, prior to the onset of cachectic syndrome. At each time point (either day 13 or day 24), all the data presented were derived from one single mouse cohort.

### Measurement of body composition

One day prior to euthanasia, the body composition of all the mice was measured by magnetic resonance-based analysis (EchoMRI 3-in-1). Unanesthetized mice were individually placed in a plastic cylinder, which were then inserted into the scanner to measure body fat, lean mass, free water mass, and total water mass.

### Muscle force testing

One day prior to euthanasia, mice were anesthetized with 2% isoflurane and placed on a platform heated to 37°C. The function of the ankle plantar flexor muscle group including the gastrocnemius (GAST), plantaris (PLA), and soleus (SOL) muscles were assessed by the non-invasive *in vivo* assay (1300A 3-in-1 Whole Animal System for Mice, Aurora Scientific). The knee joint of the right leg was immobilized by the knee clamp, and the foot was taped to the foot petal. Needle electrodes were placed superficially under the skin overlaying the GAST, and the muscle was stimulated after adjusting to optimal current to measure maximum isometric torque. To obtain the force-frequency relationship, the muscle was then stimulated at increasing frequencies (10, 25, 50, 80, 100, 150, 250, and 300 Hz) with a 1-minute rest between each stimulation. The mice were allowed to recover overnight after the *in vivo* muscle force test. On the day of euthanasia, mice were put under isoflurane anesthesia, and the distal part of the tibialis anterior (TA) muscle was isolated through dissection. During the *in-situ* force test, the knee joint was pinned to the limb plate, and the distal tendon of TA was tied to the lever arm of the force-test apparatus by a 4/0 silk suture (FST, 18020-40). The TA muscle was stimulated with the platinum needle electrodes inserted within the overlying fascia. Current and muscle length were adjusted to obtain the maximum isometric twitch force (*P*_t_) at optimal muscle length (*L*_o_). PBS was continuously applied to the exposed TA muscle. The muscle was then stimulated at a frequency of 150 Hz while held at *L*_o_ to obtain the maximum isometric tetanic force (*P*_o_). Then, TA muscle was stimulated at increasing frequencies (10, 25, 50, 80, 100, 150, 250, and 300 Hz) with a 1-minute rest period between each contraction. At the conclusion of testing *L*_o_ was measured using digital calipers, and TA optimum fiber length (L_f_) was determined by multiplying *L*_o_ by the TA muscle *L*_f_/*L*_o_ ratio of 0.6 (47). The physiological cross-sectional area (CSA) of the TA muscle was calculated as muscle mass divided by the product of *L*_f_ and 1.06 mg/mm^3^ (48). TA-specific tetanus force *sP*_o_ was calculated as *P*_o_/muscle CSA.

### Muscle tissue collection

After *in situ* force testing of the TA muscle, all the animals were euthanized via cervical dislocation and bilateral pneumothorax while under isoflurane anesthesia, and hindlimb muscles including GAST, PLA, SOL, TA, and extensor digitorum longus (EDL) from both legs were dissected and weighed. Whole muscles from the left leg and the proximal half of the right leg muscles were snap frozen in liquid nitrogen for RNA and protein extraction, and the distal half of the muscles were embedded in Optimal Cutting Temperature (OCT) compound and frozen in isopentane cooled on liquid nitrogen. All the samples were stored in –80°C freezer until downstream analysis.

### Immunocytochemistry

To assess cell morphology in the co-culture model, C2C12 myotubes were fixed in 4% paraformaldehyde (PFA, Electron Microscopy Sciences, 15710) for 30 minutes at 4°C. The cells were then permeabilized with 0.1% Triton X-100 for 30 minutes and blocked in 1% bovine serum albumin (BSA, Sigma-Aldrich A3294-50G) for 1 hour at room temperature, followed by overnight incubation with primary antibodies against sarcomeric myosin (MF20c, DSHB, 1:100) and myogenin (F5Dc, DSHB, 1:100) at 4°C. The following day, C2C12 cells were washed in PBS and then incubated with secondary antibodies including Goat Anti-Mouse IgG1 Alexa Fluor 568 (Invitrogen A-21124, Thermo Fisher Scientific, 1:500) and Goat Anti-Mouse IgG2b Alexa Fluor 647 (Invitrogen, Thermo Fisher Scientific A-21242, 1:500). DAPI (Invitrogen, Thermo Fisher Scientific D21490, 2 μg/mL) was used to counterstain cell nuclei. C2C12 myotubes were visualized using an automated fluorescence microscope (Echo Revolution) operating in inverted configuration with a 10× objective. Nine images were automatically captured from the same predetermined positions in each culture well to avoid investigator bias.

### Immunohistochemistry

Serial cross-sections (10 μm) were cut from the midbelly of OCT-embedded SOL and EDL muscles in a cryostat (Leica CM1950) at –20°C, mounted onto SuperFrost Plus slides, and air-dried. Unfixed slides were used for Hematoxylin & Eosin (H & E) and fiber type staining. For immune cell staining, slides were fixed in acetone at –20°C for 10 minutes. All the slides used for immunostaining were blocked in M.O.M immunoglobulin blocking reagent (Vector Laboratories, BMK-2202) for 1 hour at room temperature and then washing in PBS before overnight incubation at 4°C with primary antibodies. The following day, slides were washed in PBS solution and incubated with secondary antibodies for 1 hour at room temperature. After washing in PBS, the slides were mounted using Fluorescence Mounting Medium (Agilent Dako, S3023) and fluorescent images were captured using Echo Revolution microscope operating in upright configuration with a 10× objective. Primary antibodies were used to stain MyHC I (DSHB, BA-D5c, 1:100), MyHC IIA (DSHB, SC-71c, 1:100), MyHC IIB (DSHB, BF-F3c, 1:100), laminin (Abcam, ab7463, 1:100 or Novus Biologicals N-600-883, 1:100), Gr-1 (Bio-Rad, MCA2387, 1:50), CD68 (Bio-Rad, MCA1957, 1:100 or Abcam, ab283654), and CD206 (Bio-Rad, MCA2235, 1:100). Secondary antibodies used for corresponding primary antibodies included Goat Anti-Rabbit IgG (H+L) Alexa Fluor 350 conjugate (Invitrogen, Thermo Fisher Scientific, A-21068), Goat Anti-Rat IgG (H+L) Alexa Fluor 555 conjugate (Invitrogen, Thermo Fisher Scientific, A-21434), Goat Anti-Mouse IgG1 Alexa Fluor 488 conjugate (Invitrogen, Thermo Fisher Scientific, A-21121), Goat Anti-Mouse IgM Alexa Fluor 555 conjugate (Invitrogen, Thermo Fisher Scientific, A-21426), Goat Anti-Mouse IgG2b Alexa Fluor 647 conjugate (Invitrogen, Thermo Fisher Scientific, A-21242), and Goat Anti-Rabbit IgG (H+L) Alexa Fluor 647 conjugate (Invitrogen, Thermo Fisher Scientific, A-21244). DAPI (Invitrogen, Thermo Fisher Scientific, D21490, 2 μg/mL) was used to counterstain cell nuclei on the immune marker slides.

### Image analysis

In the co-culture model, overall cell density, myotube area, percentage of myogenin^+^ cells, and myotube diameter were analyzed using ImageJ/FIJI as quantitative indices for myotube differentiation and growth. To determine mean myotube diameter, the ten largest myosin-positive cells from each image were manually measured at their widest uniform point perpendicular to the length of the cell using the straight-line selection tool. For branched myotubes, each branch was measured as a single myotube, and the region where the branches join excluded from the analysis. At least six non-overlapping images were analyzed from each independent cell culture well (biological replicate). Total cell number (DAPI^+^ cells/mm^2^), percentage of myogenin^+^ cells (myogenin^+^/DAPI^+^ cells) (%), and percentage of myosin-positive area (myotube area) (%) per field of view were quantified using a custom automated in-house plugin for ImageJ/FIJI. Muscle fiber typing images were analyzed and quantified using MuscleJ 1.0.2 with minor modifications (49). Specifically, EDL and SOL samples were analyzed with customized fiber type detection sensitivity thresholds of standard deviations (SD) from the mean of SD=1 and SD=0.5, respectively. Immune cells including CD68^+^, CD206^+^, and Gr-1^+^ cells were manually counted throughout the entire EDL or SOL cross section, which was then normalized to total tissue area as determined by MuscleJ 2.0 (50). For the analysis of immune infiltration, fiber size, and fiber type composition, tissue samples from all mice (Ctrl=6, CT26=7, C26=7) were included, and the full area of the entire muscle cross section was included in all analysis.

### RNA extraction and RT-qPCR

Tissue RNA from the proximal half of snap-frozen GAST muscles (∼25 mg) or a uniform piece of snap-frozen isolated tumor tissue (∼30 mg) was extracted using TRIzol Reagent (Invitrogen, ThermoFisher Scientific, 15596018). GAST tissue or tumor samples were homogenized using in the Mixer Mill MM 400 (Retsch) bead homogenizer for 10 minutes in safe-lock microtubes containing zirconium beads. Then, bromochloropropane was used for supernatant phase separation prior to RNA isolation with assistance with Direct-zol^TM^ RNA MiniPrep Plus kit (Zymo Research, R2072), and on-column DNase treatment. RNA concentration and purity was assessed using the Nanodrop 2000c UV-Vis Spectrophotometer (Thermo Scientific). Following cDNA synthesis using High-Capacity cDNA Reverse Transcription Kit (Applied Biosystems, Thermo Fisher Scientific, 4368813), the level of mRNA was measured by reverse transcription quantitative polymerase chain reaction (RT-qPCR) using SYBR^TM^ Green PCR Master Mix (Applied Biosystems, Thermo Fisher Scientific, 4312704). RT-qPCR plates were run using the CFX Opus 96 Real-Time PCR System (Bio-Rad Laboratories). Several reference genes were tested for expression stability. The mRNA expression of Valosin-containing protein (VCP, *Vcp*), ER membrane protein complex subunit 7 (EMC7, *Emc7*), and GAPDH (*Gapdh*) were identified as the least variable and were used as reference genes. The geometric mean of these three reference genes was used for normalization. The sequences of all the primers used are shown in **Table 1**. RT-PCR data were analyzed using the 2^−ΔΔCT^ method.

**Table 1.**
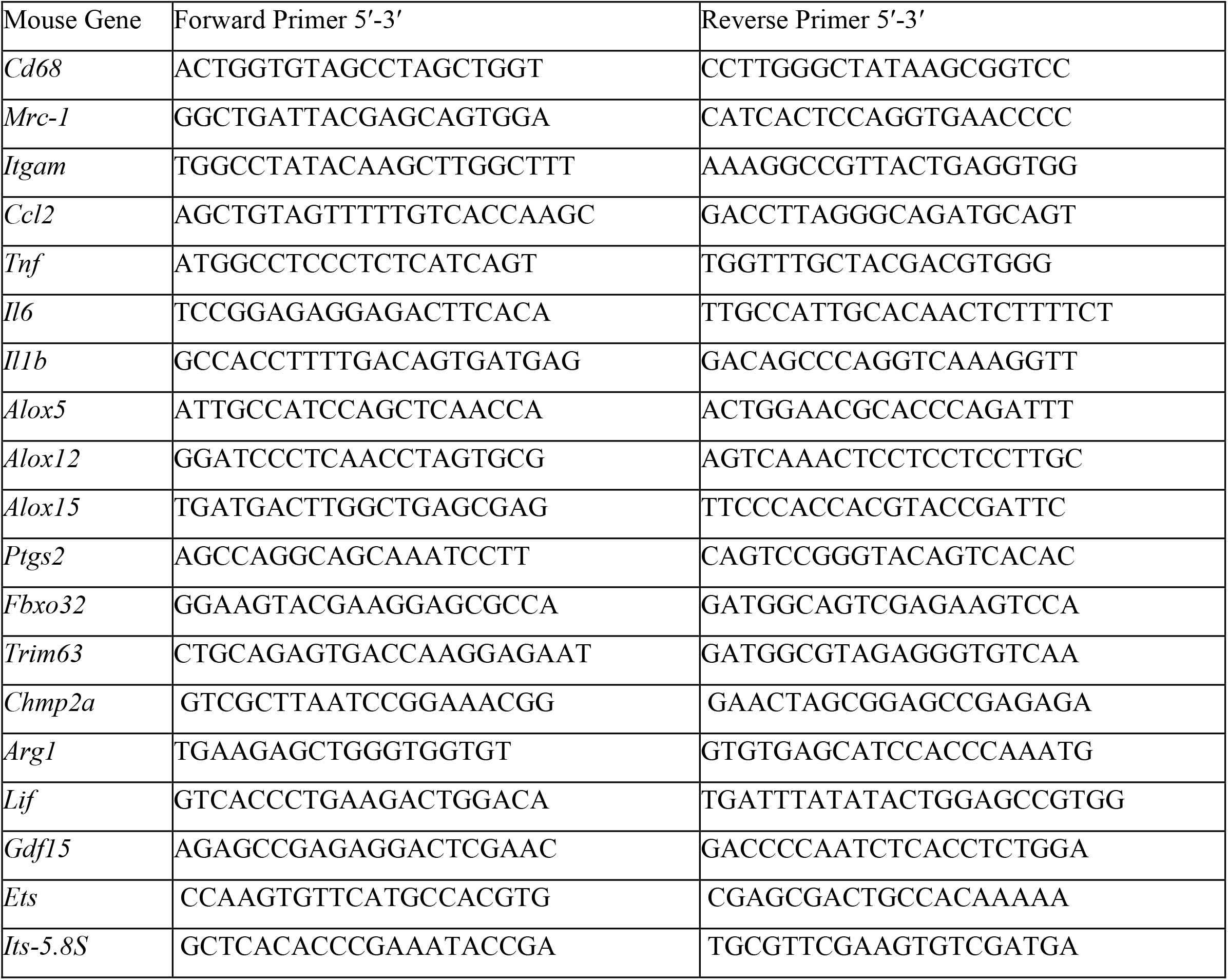
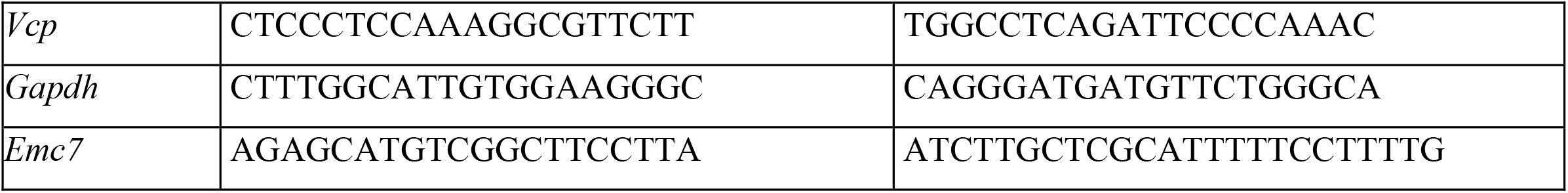
Primer sequences for RT-qPCR.

### Protein extraction and western blot

Frozen GAST tissue from the left leg (∼40 mg) was homogenized in ice-cold 1× RIPA buffer (MilliporeSigma, 20188, 15 μL/mg tissue) with 1× Halt Protease and Phosphatase Inhibitor Cocktail (Thermo Fisher Scientific, 78442) using a Fisherbrand Bead Mill 4 mini homogenizer (Thermo Fisher Scientific, 15-340-164) at 5 m/s for 60 seconds. Tissue homogenates were incubated with mild agitation at 4°C for 1 hour. After centrifugation at 13,000 × *g* for 10 minutes at 4°C, the supernatant containing protein was collected. Pierce bicinchoninic acid (BCA) protein assay kit (Thermo Fisher Scientific, PI23225) was used to determine protein concentration in each sample, and protein samples were diluted to 2 μg/μL in 1 × Laemmli buffer (Bio-rad, 1610737). The protein samples were boiled for 5 min, and an equal volume of protein (20 μg) was loaded to a 12% sodium dodecyl sulfate-polyacrylamide gel electrophoresis (SDS-PAGE). After separation by gel electrophoresis, the protein samples were transferred to a nitrocellulose membrane using a Trans-Blot Turbo Transfer System (Bio-Rad, 1704150). Membranes were blocked in 5% skim milk powder in Tris buffered saline with 0.1% Tween 20 (TBST) for 1 hour at room temperature. Then, membranes were incubated with primary antibodies diluted in 5% bovine serum albumin (BSA) in TBST overnight at 4 °C with gentle agitation. Primary antibodies tested included Rabbit Anti-15-LOX-1 (Abcam, ab244205, 1:1,000), Rabbit Anti-S6 Ribosomal Protein (Cell Signaling Technology, 2217, 1:1,000), Rabbit Anti-Phospho-S6 Ribosomal Protein (Ser235/236) (Cell Signaling Technology, 4858, 1:1,000), Rabbit Anti-Phospho-S6 Ribosomal Protein (Ser240/244) (Cell Signaling Technology, 5364, 1:1,000), Rabbit Anti-p70 S6 Kinase (Cell Signaling Technology, 2708, 1:1,000), Rabbit Anti-Phospho-p70 S6 Kinase (Thr421/Ser424) (Cell Signaling Technology, 9204, 1:1,000), Rabbit Anti-Phospho-p70 S6 Kinase (Thr389) (Cell Signaling Technology, 9234, 1:1,000), Rabbit Anti-Phospho-SAPK/JNK (Thr183/Tyr185) (81E11) (Cell Signaling Technology, 4668, 1:1000), Rabbit Anti-SAPK/JNK (Cell Signaling Technology, 9252, 1:1000), Rabbit Anti-Phospho-Akt (Ser473) (D9E) (Cell Signaling Technology, 4060, 1:1000), Rabbit Anti-Phospho-Akt (Thr308) (D25E6) (Cell Signaling Technology, 13038, 1:1000), Rabbit Anti-Akt (pan) (Cell Signaling Technology, 4691, 1:1000), Rabbit Anti-p44/42 MAPK (Erk1/2) (Cell Signaling Technology, 4695, 1:1,000), and Rabbit Anti-Phospho-p44/42 MAPK (Erk1/2) (Thr202/Tyr204) (Cell Signaling Technology, 4370, 1:1,000). Mouse Anti-GAPDH was used to probe for the loading control (Santa Cruz Technology, sc-32233, 1:1,000). The following day, membranes were washed in TBST for 5 minutes × 3 times and probed with a Goat Anti-Rabbit IgG (H + L) Horseradish Peroxidase (HRP) conjugated secondary antibody (Jackson Labs, 111-035-144, 1:10,000) or Goat Anti-Mouse IgG (H + L) Horseradish Peroxidase (HRP) conjugated secondary antibody (Jackson Labs, 115-035-003, 1:10,000) diluted in 5% skim milk in TBST for 1 hour at room temperature. Protein bands were visualized using Clarity Western enhanced chemiluminescent (ECL) substrate (Bio-Rad, 1705060), and chemiluminescent signals were captured by a ChemiDoc Imaging System (Bio-Rad, 12003153). Densitometry analysis was performed using Image Lab 6.1 software (Bio-Rad).

### Statistical analysis

Data are presented as the group mean ± SEM, with raw data from each biological replicate overlaid on bar graphs. Statistical analysis was performed in GraphPad Prism 10. In mouse experiments with one independent variable and three groups (i.e., Ctrl vs. CT26 vs. C26), differences between groups were tested using a one-way ANOVA. Following a significant one-way ANOVA, post hoc tests were performed using Fisher’s least significant difference (LSD) test, which controls the family-wise Type I error rate in the unique scenario of three experimental groups while maintaining statistical power. For mouse experiments with two independent variables (e.g., stimulation frequency and cell type, or time and cell type), data were analyzed using a two-way repeated-measures ANOVA. Following a significant main effect or interaction, pairwise comparisons among the three groups (Ctrl vs. CT26 vs. C26) within each frequency or time point were performed using Fisher’s LSD post hoc test. For *in vitro* experiments with four experimental groups (C2C12 control, low, middle, and high carcinoma cell densities), data were analyzed with a two-way ANOVA followed by Holm–Šidák post hoc tests. *P* ≤ 0.05 was used to determine statistical significance. Statistical outliers were identified by Grubbs’ test (α = 0.05), with at most one outlier removed from each group.

## RESULTS

### C26 and CT26 Cells Exhibit Similar Potency in Inducing *In Vitro* Muscle Atrophy

To investigate the effect of soluble factors secreted by C26 and CT26 carcinoma cells to induce skeletal muscle cell wasting *in vitro*, we co-cultured mature C2C12 myotubes with increasing densities of C26 or CT26 cells (Low: 4.17 × 10^4^/cm^2^, Middle: 8.3 × 10^4^/cm^2^, and High: 1.67 × 10^5^/cm^2^) to allow for continuous exchange of muscle- and cancer-secreted factors (**Figure 1A**). C2C12 myotubes co-cultured with C2C12 myoblasts served as the control group. Immunocytochemistry analysis showed that 72-hours of co-culture with either C26 or CT26 cells induced significant atrophy of C2C12 myotubes (**Figure 1B**). Both C26 and CT26 cells led to a marked decrease in C2C12 myotube diameter (**Figure 1C**). In addition, C26 and CT26 cells both reduced overall myotube area expressed as a percentage of the total field of view area at middle and high densities (**Figure 1D**). While CT26 cells did not influence myogenin expression, C2C12 myotubes co-cultured with C26 cells also had a reduced proportion of myogenin^+^ cells at low and middle cancer cell co-culture densities (**Figure 1E**). There was no effect of carcinoma cell density on C2C12 myotube viability as indicated by the total number of DAPI^+^ cells/mm^2^ in the base plates (**Figure 1F**). However, at middle density, CT26 cell co-culture did result in a lower viability of the C2C12 myotube cultures when compared to C26 cells (**Figure 1F**). Overall, these data show that both CT26 and C26 carcinoma cells induce robust *in vitro* skeletal muscle cell wasting.

**Figure 1.**
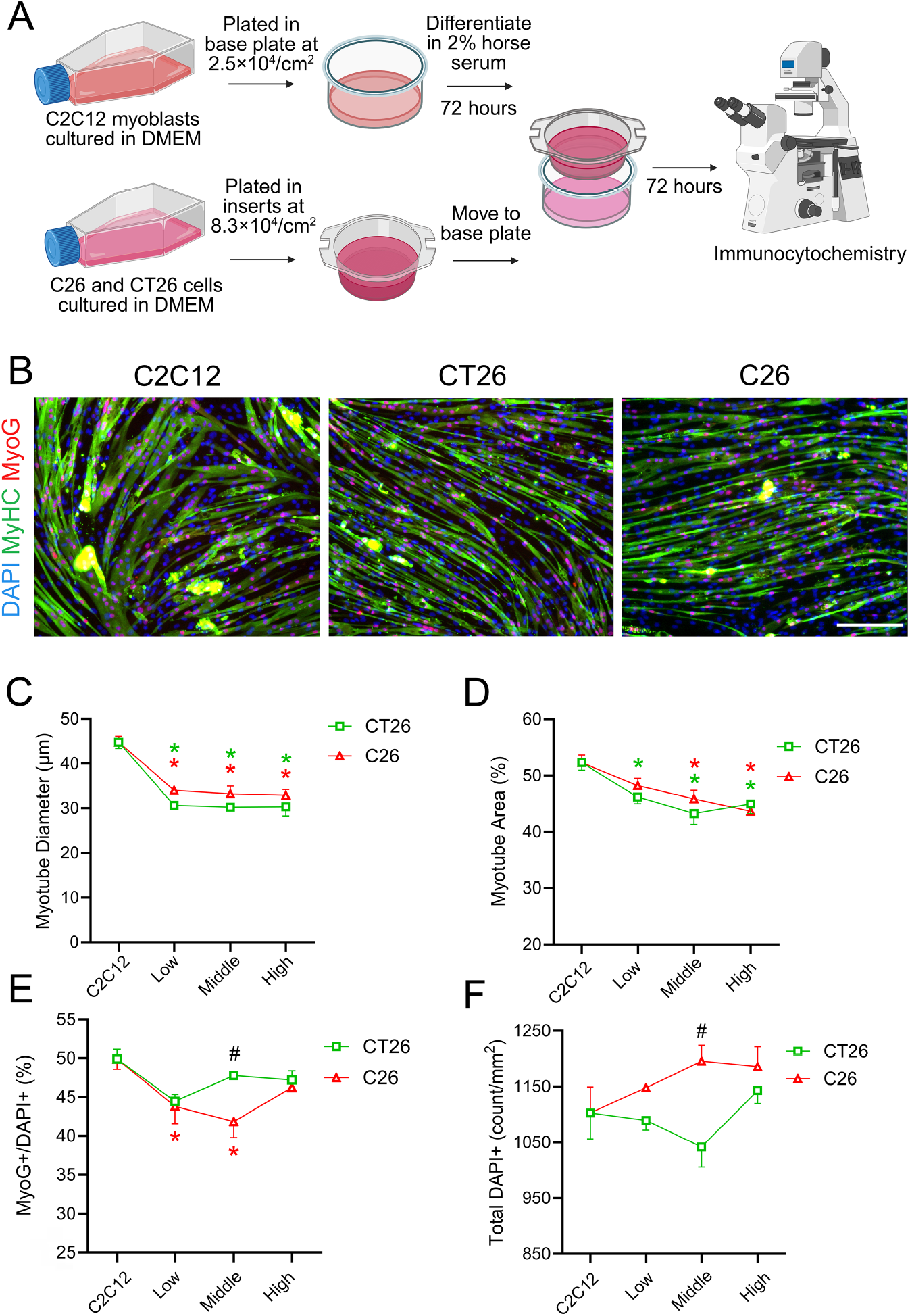
CT26 and C26 carcinoma cells induced *in vitro* muscle atrophy in a cancer-muscle co-culture model. **(A)** Subconfluent C2C12 myoblasts cultured in growth media consisting of DMEM containing 10% fetal bovine serum were plated in the 24-well base plate at a cell density of 2.5 × 10^4^/cm^2^. C2C12 myoblasts were allowed to proliferate for four days and were then induced to form myotubes by switching to differentiation media consisting of DMEM containing 2% horse serum for 72 hours. On the same day as induction of differentiation, C26 cells and CT26 cells were plated in growth media in co-culture inserts placed in empty base plates at increasing densities (4.17 × 10^4^/cm^2^, 8.3 × 10^4^/cm^2^, and 1.67 × 10^5^/cm^2^) and allowed to proliferate for 72 hours. C2C12 myoblasts were plated in the inserts at 8.3 × 10^4^/cm^2^ as a negative co-culture control. On day 3 of myogenic differentiation, the co-culture inserts containing confluent C26 cells, CT26 cells, or C2C12 myoblasts were moved to the lower compartments of the base plate containing mature C2C12 myotubes. Fresh differentiation media was added to both the inserts and lower compartments when establishing the co-cultures. **(B)** Following 72 hours of co-culture the C2C12 myotubes in the base plate were fixed with 4% paraformaldehyde (PFA) and stained for myosin heavy chain (MyHC, green) and myogenin (MyoG, red). DAPI was used to counterstain the nuclei (blue). Scale bar is set at 200 µm. Myotube diameter **(C)**, myotube area expressed as a percentage of total field of view area **(D)**, MyoG^+^/total DAPI^+^ percentage **(E)**, and total DAPI^+^ cell density **(F)** was quantified to assess potential atrophic effect of C26 cells and CT26 cells on C2C12 myotubes. Data are presented as the mean ± SEM of three independent co-culture wells (*n* = 3). Statistical significance was determined by two-way ANOVA followed by Holm–Šidák post hoc tests. \**p* < 0.05 for pairwise comparison to C2C12 insert control, #*p* < 0.05 for comparison between C26 and CT26 cells.

### C26 Tumors Induce a Larger Loss in Body Weight, Lean Mass, and Fat Mass

To study the potential of C26 and CT26 tumors to induce cancer cachexia *in vivo*, three groups of female BALB/c mice with the same initial body weight were subcutaneously injected in the right flank with saline (Ctrl), C26 cells (5 × 10^5^ cells/mouse), or CT26 cells (5 × 10^5^ cells/mouse) (**Figure 2A**). Tumors formed by C26 and CT26 cells became palpable and measurable 5 days after inoculation. Although the planned final terminal tissue collection was scheduled for day 24 post inoculation, some tumor-bearing mice began to reach humane endpoints starting on day 19. Consequently, differences in the exact duration of longitudinal measurements reflect animal attrition between day 19 and day 24 of the study. Notably, mice inoculated with CT26 cells developed larger subcutaneous tumors when compared to those inoculated with C26 cells as indicated by both more rapid estimated growing tumor volume (**Figure 2B)** and final terminal tumor mass (**Figure 2C)**. However, both CT26 and C26 tumor-bearing mice ultimately had a similar probability of survival (**Figure 2D**). Both groups of tumor-bearing mice showed a compensatory increase in food intake during the early stages of tumor growth, but C26 mice started to consume less food than CT26 mice on day 14 following tumor cell inoculation (**Figure 2E**). Terminal EchoMRI analysis of body composition revealed that both C26 and CT26 tumors induced fat wasting, but C26 tumor-bearing mice experienced greater loss of fat mass than CT26 mice (**Figure 2F**). On the other hand, only C26 tumors resulted in significantly lower tumor-free lean mass than healthy control mice at endpoint (**Figure 2G**). In addition, while C26 tumor-bearing mice experienced a significant loss of total body weight starting day 13 post inoculation, CT26 tumor-bearing mice did not exhibit any such loss in body weight (**Figure 2H**). Both C26 and CT26 tumors led to a marked reduction in estimated tumor-free body weight, although C26 tumors exhibited both an earlier onset and greater final magnitude of weight loss than CT26 mice (**Figure 2I**). CT26 and C26 tumor-bearing mice both had significantly increased endpoint spleen mass, demonstrating high systemic inflammation induced by tumor progression (**Figure 2J**). Both C26 and CT26 tumors caused a significant loss of GAST and TA muscle mass when expressed relative to starting body weight, although C26 tumors resulted in a relatively greater amount of muscle wasting than CT26 mice (**Figure 2K).** A similar response was seen for absolute terminal GAST and TA muscle mass (**Figure S1A**). However, in smaller hindlimb muscle groups including the PLA, EDL, and SOL, only C26 tumors induced a significant loss of muscle mass relative to starting body weight (**Figure 2K)** and absolute muscle mass (**Figure S1A**) when compared to healthy controls.

**Figure 2.**
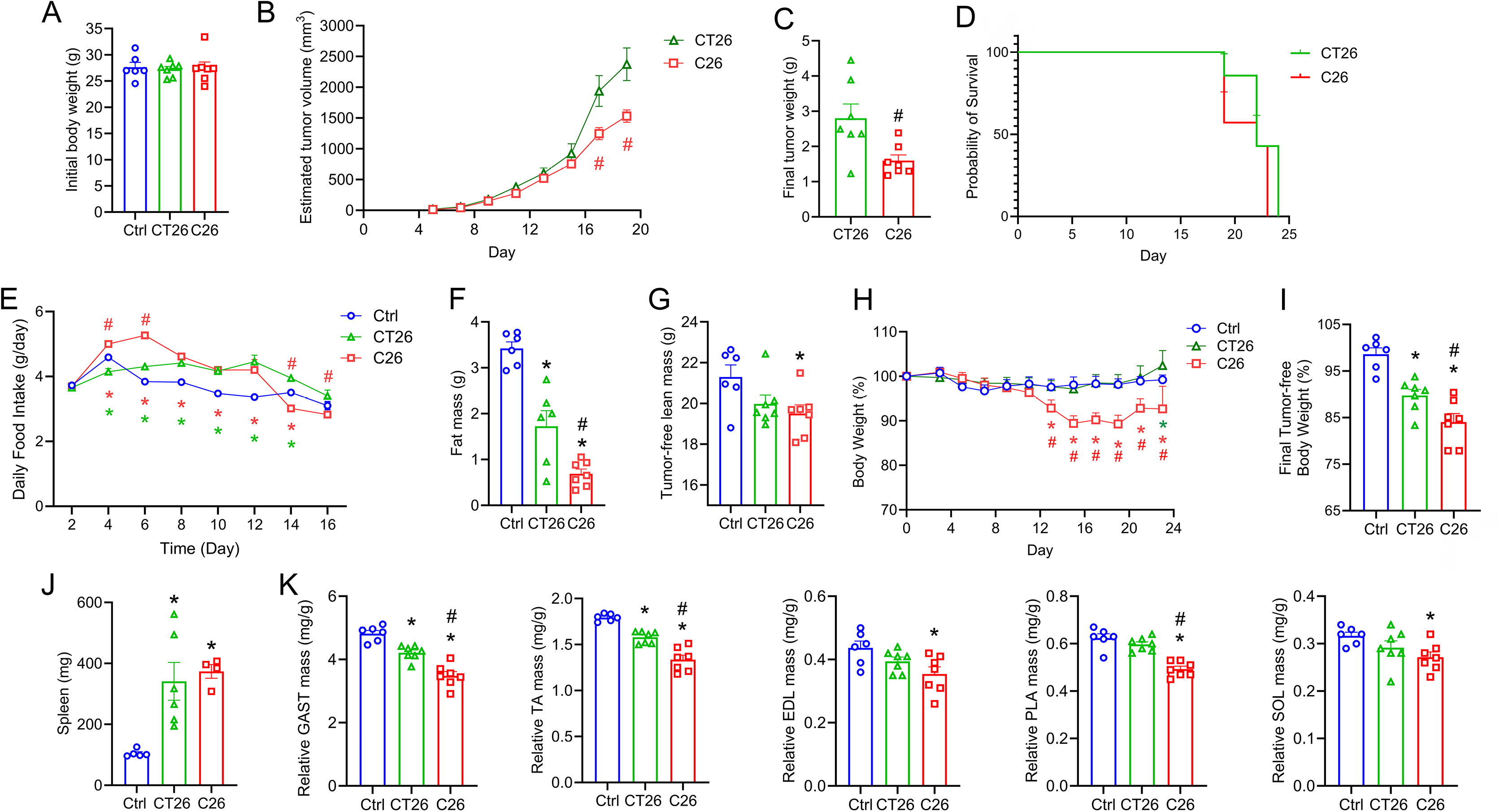
C26 and CT26 tumor inoculation induced changes in body weight and body composition. **(A)** Female BALB/c mice were randomized to three groups with similar initial body weights and injected subcutaneously into the right flank with saline (Ctrl) (*n* = 6), 5 × 10^4^ CT26 cells (*n* = 7), or 5 × 10^4^ C26 cells (*n* = 7). **(B)** Tumor diameters were measured every other day, and estimated tumor volume was calculated as 0.52 × length × width^2^. **(C)** Final tumor weight was determined after dissection on the day of euthanasia. **(D)** Probability of survival curve was plotted based on mice being euthanized after meeting the humane endpoint criteria or reaching 24 days post inoculation. **(E)** Daily food intake for each group was estimated by measuring the weight of food provided and left over pellets every 48 hours. **(F)** One day prior to euthanasia, EchoMRI was used to assess body composition including lean and fat mass. **(G)** Tumor-free lean mass was calculated as absolute lean mass minus endpoint tumor mass. **(H)** Total body weight was assessed every other day to calculate percentage change of absolute body weight. **(I)** Final tumor-free body weight was calculated as final body weight minus final tumor weight. **(J)** The mass of spleen was recorded after euthanasia. **(K)** Relative mass of major hindlimb skeletal muscles after normalization to initial body weight including the gastrocnemius (GAST), tibialis anterior (TA), extensor digitorum longus (EDL), plantaris (PLA), and soleus (SOL). Data are presented as the mean ± SEM, with dot plots showing raw data obtained from each individual mouse (biological replicates). Statistical significance was determined by one-way ANOVA (panels A, F, G, I, J, and K) or two-way ANOVA (panels B, E, and H) followed by Fisher’s LSD post hoc tests. Terminal tumor weight (panel C) was compared between the CT26 and C26 groups by two-tailed unpaired t-tests. \**p* < 0.05 for difference compared to healthy controls, #*p* < 0.05 for difference between C26 and CT26 tumor-bearing mice.

### C26 Tumors Induce Greater Decrements in Muscle Strength Compared to CT26

Next, we investigated the influence of C26 and CT26 tumors on skeletal muscle function. To test whether tumor progression affects GAST strength, all mice were tested for the maximal plantar flexion torque *in vivo* one day prior to euthanasia. CT26 tumors decreased the single twitch torque of GAST muscle (**Figure 3A**) and had a trend to reduce the absolute peak torque during tetanus (**Figure 3B**). On the other hand, while C26 tumors did not result in significantly reduced twitch torque (**Figure 3A**), they induced a greater reduction of maximal tetanic torque compared to CT26 tumor-bearing mice (**Figure 3B**). Similarly, C26 tumor-bearing mice exhibited a lower relative tetanic torque when normalized to final terminal body weight when compared to either the CT26 or healthy control group (**Figure 3C**). Moreover, C26 tumors led to a decrease in maximal isometric torque when stimulated by medium to high frequencies (80-300 Hz), while no difference was detected in CT26 tumor-bearing mice (**Figure 3D**).

**Figure 3.**
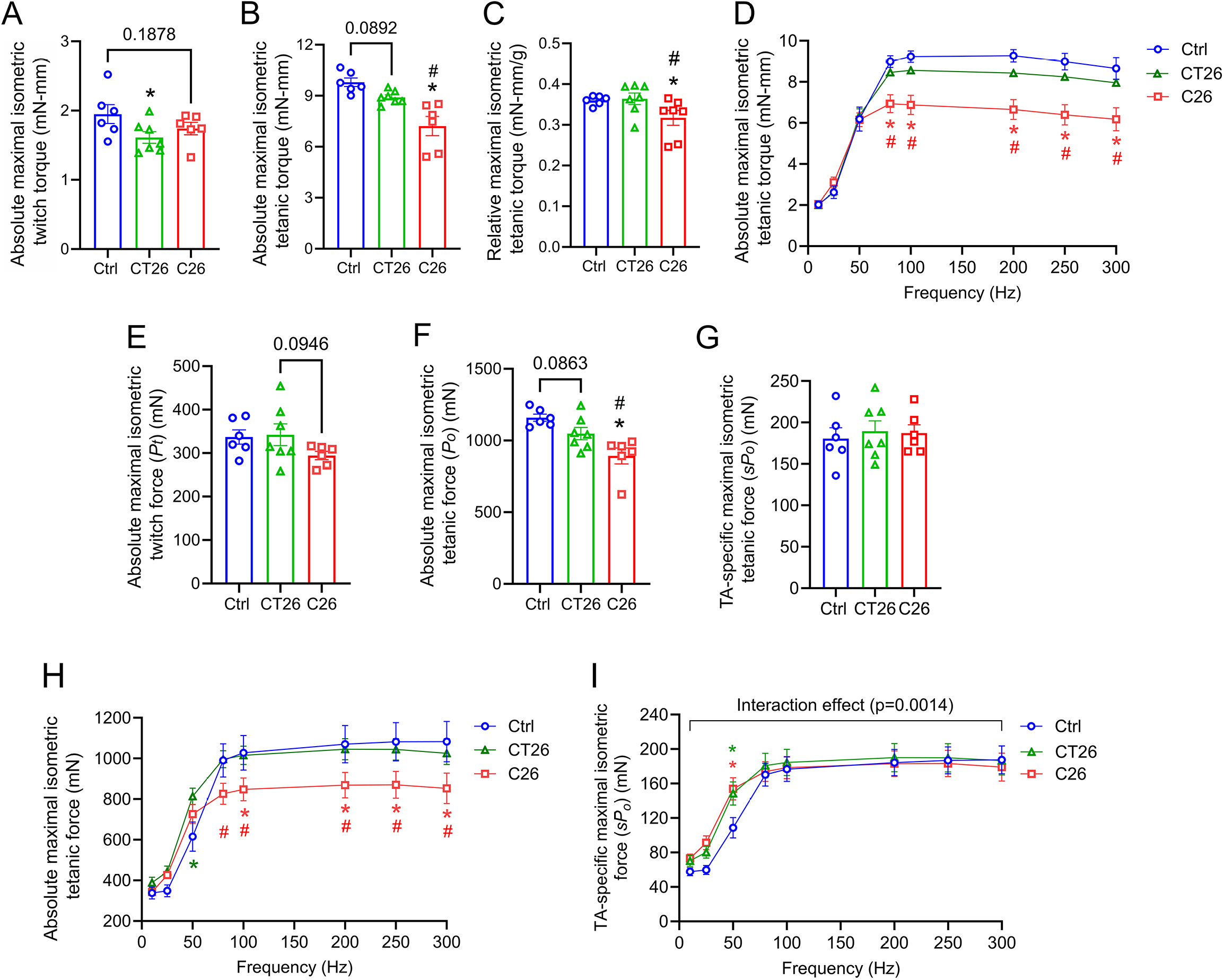
C26 and CT26 tumors differentially impacted the muscle mass and force output of BALB/c mice. To determine the force output of the gastrocnemius muscle (GAST), absolute maximal isometric twitch torque **(A)** and maximal isometric tetanic torque **(B)** were measured by *in vivo* force test one day prior to euthanasia. **(C)** Relative tetanic torque after normalization to terminal body weights. **(D)** The force-frequency relationship of the GAST muscle was generated by muscle stimulation at increasing electric frequencies (10, 25, 50, 80, 100, 150, 250, 300 Hz) with a 1-minute rest between contractions. After overnight recovery, the force output of tibialis anterior (TA) muscle was assessed by maximal isometric twitch force **(E)** and maximal isometric tetanic force **(F). (G)** Maximal TA-specific tetanus force was calculated following normalization to muscle cross-sectional area (CSA). **(H)** The force-frequency curve of the TA was generated like that of the GAST. **(I)** Specific force-frequency curves of the TA are also shown following normalization of absolute strength to muscle CSA. Data are presented as the mean ± SEM, with dot plots showing raw data obtained from each individual mouse (biological replicates). Statistical significance was determined by one-way ANOVA (panels A, B, C, E, F, and G) or two-way ANOVA (panels D, H, and I) followed by Fisher’s LSD post hoc tests. \**p* < 0.05 for difference compared to healthy controls, #*p* < 0.05 for difference between C26 and CT26 tumor-bearing mice.

To assess the potential change in the contractile function of the TA, the hindlimb muscle primarily responsible for ankle dorsiflexion, we performed the *in-situ* force test on TA muscle after overnight recovery following the GAST force test. C26 tumor-bearing mice tended to have a reduced isometric twitch force (*p* = 0.096) (**Figure 3E**). CT26 tumors tended to reduce the maximal tetanic force of the TA (*p* = 0.083), but C26 tumors resulted in a considerably greater reduction in TA tetanus force when compared to the CT26 group (**Figure 3F**). When TA muscle strength was normalized to physiological muscle cross-sectional area (CSA), no difference was observed in TA-specific tetanus force produced by the healthy controls and the C26 or CT26 tumor-bearing mice (**Figure 3G**). When the TA was stimulated at increasing frequencies, C26 tumor-bearing mice displayed lower absolute isometric force production when compared to healthy controls or CT26 mice at stimulation frequencies between 80 and 300 Hz (**Figure 3H**). However, analysis of the TA specific force frequency relationship after normalization of muscle CSA revealed that both C26 and CT26 tumor-bearing mice generated greater relative strength than healthy control mice, but only at a low stimulation frequency of 50 Hz (**Figure 3I**).

### C26 and CT26 Tumors Result in Distinct Changes in Skeletal Muscles Fiber Morphology

To better characterize how C26 and CT26 tumors influence different muscle types, we investigated the changes in muscle morphology using the EDL and SOL as representative types of fast-twitch and slow-twitch muscles, respectively (**Figure 4A-C**). Immunohistochemistry staining showed that C26 tumors tended to decrease the mean myofiber CSA of the EDL (*p* = 0.072) (**Figure 4D**) and significantly reduced the mean myofiber CSA of the SOL (**Figure 4H**). CT26 also showed a statistical trend to decrease myofiber CSA of the SOL (*p* = 0.089) (**Figure 4H**), but not the EDL (**Figure 4D**). The fiber size frequency distribution of the EDL (**Figure 4E**) and SOL (**Figure 4I**) further showed that C26 tumor-bearing mice exhibited a shift towards relatively smaller myofibers. We did not observe any changes in the percentage fiber type composition of either the EDL (**Figure 4F**) or SOL (**Figure 4J**). However, the atrophic effect of C26 tumors in the EDL was predominantly caused by fast-twitch type IIB fiber atrophy, while the type I fibers rather tended to be larger in EDL muscles of C26 tumor-bearing mice (**Figure 4G**). Intriguingly, the decrease in muscle fiber size in the SOL of C26 tumor-bearing mice was mainly driven by the atrophy of slow-twitch type I muscle fibers (**Figure 4K**). No significant difference was detected in CT26 tumor-bearing mice for fiber type-specific CSA in either the EDL (**Figure 4G**) or SOL (**Figure 4K**). Overall, C26 tumor development resulted in a higher level of muscle atrophy in both the slow-twitch SOL and the fast-twitch EDL.

**Figure 4.**
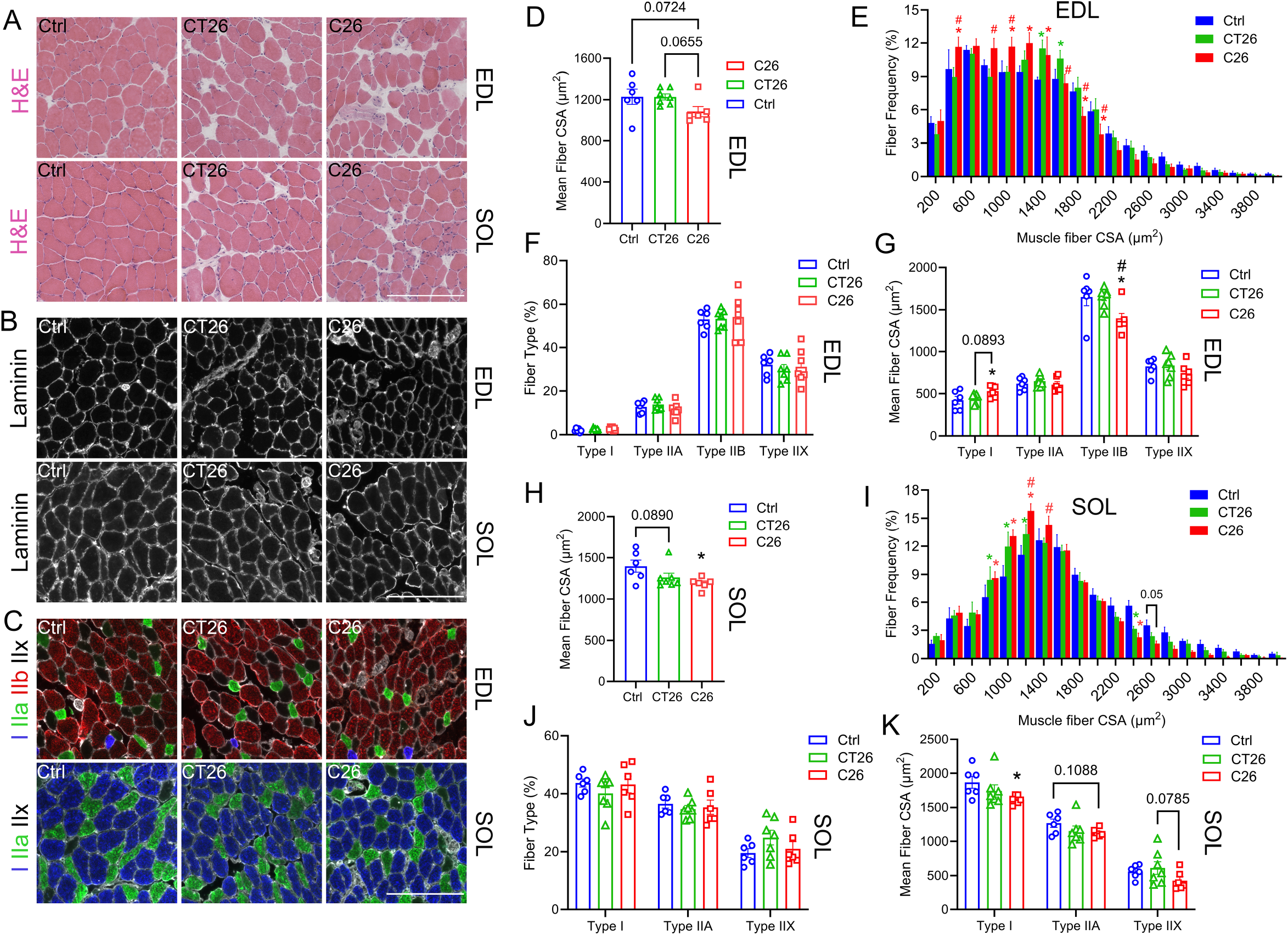
C26 and CT26 tumors resulted in morphological changes in fast-twitch and slow-twitch muscles. **(A)** Overall tissue morphology of extensor digitorum longus (EDL) and soleus (SOL) from healthy controls, C26 tumor-bearing mice, and CT26 tumor-bearing mice were visualized by hematoxylin and eosin (H&E) staining. **(B)** The muscle basement membranes of EDL and SOL were visualized by immunostaining for laminin (white). **(C)** Different muscle fiber types were identified in EDL and SOL by immunofluorescence. Myosin heavy chain I (MyHC I) fibers were stained by BA-D5c (blue), MyHC IIA was stained by SC-71c (green), and MHC IIB was stained by BF-F3c (red). Scale bar is set at 200 µm. Fiber morphology of EDL muscles was assessed by mean cross-sectional area (CSA) **(D)**, fiber size distribution frequency **(E)**, fiber type percentage **(F)**, and fiber type-specific mean CSA **(G)**. SOL fiber morphology was also determined by mean CSA **(H)**, fiber size distribution frequency **(I)**, fiber type percentage **(J)**, and fiber type-specific mean CSA **(K).** Data are presented as the mean ± SEM, with dot plots showing raw data obtained from each individual mouse (biological replicates). Statistical significance was determined by one-way ANOVA followed by Fisher’s LSD tests. \**p* < 0.05 for difference compared to healthy controls, #*p* < 0.05 for difference between C26 and CT26 tumor-bearing mice.

### Differential Intramuscular Immune Responses of C26 and CT26 Tumor-bearing Mice

To determine potential differences in skeletal muscle immune cell infiltration during cancer cachexia, total macrophages (MФ) (CD68^+^ cells), M2-like MФ (CD68^+^CD206^+^ cells), and polymorphonuclear neutrophils (PMNs, Gr-1^+^ cells) were stained in EDL (**Figure 5A and 5B)** and SOL (**Figure 5F and 5G**). In the EDL, C26 tumor-bearing mice showed higher intramuscular numbers than either healthy controls or the CT26 group of total MФ (**Figure 5C**), M2-like MФ (**Figure 5D**), and PMNs (**Figure 5E**). In the SOL, both C26 and CT26 tumors resulted in greater intramuscular infiltration than healthy controls of total MФ (**Figure 5H)** and M2-like MФ (**Figure 5I**), although C26 tumor-bearing mice showed a higher level of MФ infiltration than the CT26 group. C26 tumor-bearing mice also showed infiltration of the SOL by PMNs, and a similar trend was observed in the CT26 group (*p* = 0.112) (**Figure 5J**).

**Figure 5.**
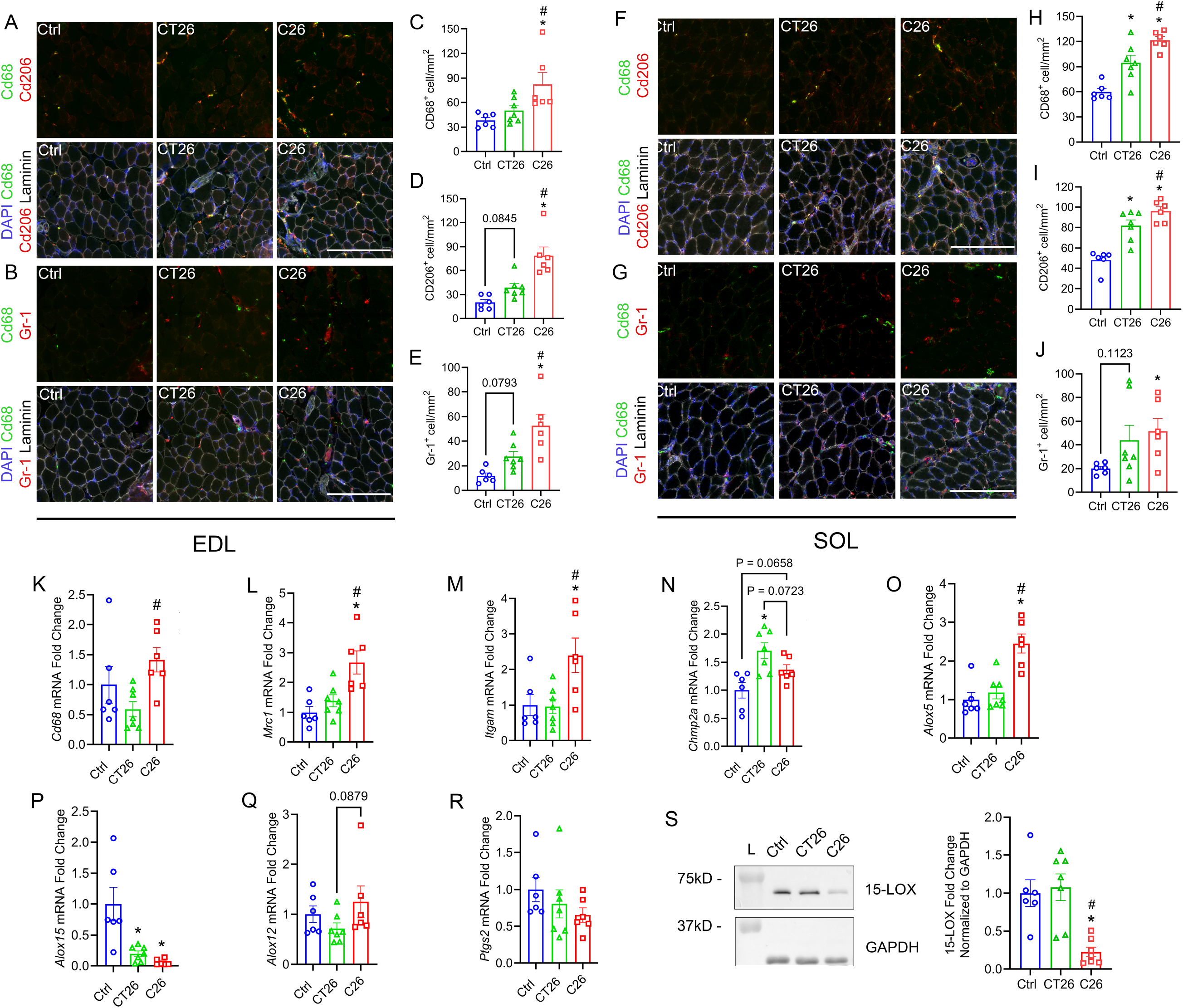
C26 tumor-bearing mice exert a higher level of intramuscular immune response than CT26 tumor-bearing mice. Sections of EDL **(A-B)** and SOL **(F-G)** were stained for immune cells including total macrophages (MΦ) (CD68^+^ cells), M2-like MΦ (CD206^+^ cells), and polymorphonuclear neutrophils (PMNs) (Gr-1^+^ cells). Intramuscular immune infiltration was assessed by the quantification of CD68^+^ cells, CD68^+^CD206^+^ cells, and Gr-1^+^ cells in EDL **(C-E)** and SOL **(H-J)**. Scale bar is set at 200 µm. The mRNA expression level of *Cd68* **(K)**, *Mrc1* **(L),** *Itgam* **(M)**, *Chmp2a* **(N)**, *Alox5* **(O)**, *Alox15* **(P),** *Alox12* **(Q),** and *Ptgs2* **(R)** was determined by real-time quantitative reverse transcription PCR (RT-qPCR). Gene expression was normalized to the geometric mean of *Vcp*, *Emc7*, and *Gapdh*. Western blot was used to examine the protein expression of leukocyte-type 12/15-LOX **(S)**. Data are presented as the mean ± SEM, with dot plots showing raw data obtained from each individual mouse (biological replicates). Statistical significance was determined by one-way ANOVA followed by Fisher’s LSD post hoc tests. \**p* < 0.05 for difference compared to healthy controls, #*p* < 0.05 for difference between C26 and CT26 tumor-bearing mice.

To further study the transcriptional regulation of the skeletal muscle inflammatory response, we measured mRNA expression of innate immune markers in the GAST muscle including CD68 (*Cd68*) (**Figure 5K**), CD206 (*Mrc1*) (**Figure 5L**), and CD11b (*Itgam*) (**Figure 5M**). While *Cd68* expression was not different between CT26 tumor-bearing mice and healthy controls, C26 tumors induced a significant increase in muscle *Cd68* expression when compared to the CT26 group (**Figure 5K**). Meanwhile, we observed a significant increase in expression of *Mrc1* (**Figure 5L)** and *Itgam* (**Figure 5M**) relative to healthy controls in C26 but not CT26-bearing mice. We also measured the expression of charged multivesicular body protein 2A (CHMP2A, *Chmp2a*), which is known to regulate autophagy and mediates the cytotoxicity of natural killer (NK) cells in tumor immunity (51, 52). CT26 tumors induced an increase in *Chmp2a* expression, and a similar trend was observed in the C26 group (*p* = 0.066) (**Figure 5N**).

We also measured GAST muscle expression of major pro-inflammatory cytokines including monocyte chemoattractant protein-1 (MCP-1, *Ccl2*) (**Figure S1B**), tumor necrosis factor-alpha (TNFα, *Tnf*) (**Figure S1C**), interleukin-6 (IL-6, *Il6*) (**Figure S1D**), and interleukin-1 beta (IL-1β, *Il1b*) (**Figure S1E**). Intriguingly, while C26 did not change the expression of *Ccl2* or *Tnf*, CT26 tumors repressed muscle expression of both *Ccl2* and *Tnf* (**Figure S1B and S1C**). There was no difference in muscle expression of *Il6* and *Il1b* in either C26 or CT26 tumor-bearing mice (**Figure S1D and S1E**). Additionally, there were no differences between groups in the expression of external transcribed spacer (ETS, *ETS*) transcription factors or internal transcribed spacer-5.8S ribosomal DNA region (ITS-5.8S, *ITS-5.8S*), which is involved in skeletal muscle ribosome biogenesis (**Figure S1F and S1G**) (53).

We then assessed the expression of key enzymes involved in the biosynthesis of lipid mediators and immune regulation, including 5-lipoxygenase (5-LOX, *Alox5*) (**Figure 5O**), leukocyte-type 12/15-LOX (12/15-LOX, *Alox15*) (**Figure 5P**), platelet-type 12-LOX (12-LOX, *Alox12*) (**Figure 5Q**), and cyclooxygenase 2 (COX-2, *Ptgs2*) (**Figure 5R**). We found that C26 tumors significantly increased muscle expression *Alox5*, which was not observed in mice bearing CT26 tumors (**Figure 5O**). Interestingly, both C26 and CT26 tumors greatly diminished the expression of *Alox15*, which plays an important role in the production of specialized pro-resolving mediators (SPMs) and the resolution of muscle inflammation (**Figure 5P**) (54, 55). Further analysis of muscle 15-LOX protein expression by western blot confirmed a robust downregulation of 15-LOX at the protein level in C26 tumor-bearing mice, but 15-LOX was not reduced in CT26 mice (**Figure 5S**). COX-2 and 12-LOX have been shown to promote CRC cachexia through activating inflammation and repressing protein synthesis pathways (32, 56). Nevertheless, muscle expressions of *Alox12* (**Figure 5Q**) and *Ptgs2* (**Figure 5R**) were not impacted by either C26 or CT26 tumors.

### Similar Mechanistic Signatures in Muscles at Early Cachexia Stages

To address whether the observed differential immune profile and lipid biosynthesis pathways were a potential underlying cause of cachexia or simply a phenotypic result of advanced cachexia stage, we further compared muscle and tumor outcomes in mice at an earlier time-point of 13 days following tumor cell inoculation. At this point neither CT26 nor C26 tumor-bearing mice had yet lost a significant amount of body weight (**Figure S2A**). There was also not yet any difference in lean or fat mass between tumor-bearing mice and healthy controls, although C26 mice did already display a lower absolute fat mass when compared to CT26 mice (**Figure S2A**). CT26 tumor-bearing mice were generally found to consume more food than C26 tumor-bearing mice during this early window following tumor cell implantation (**Figure S2B**). Despite this, no differences were yet observed for estimated tumor volume or final tumor mass between the CT26 and C26 groups (**Figure S2C**). Nevertheless, spleen mass was already found to be markedly increased in the C26 group (**Figure S2D**). Despite this, neither CT26 nor C26 tumor-bearing mice yet differed from healthy control mice for the mass of the TA, EDL, SOL, PLA, and GAST when expressed relative to starting body weight (**Figure S2E**). Force output data also indicated that muscle strength did not yet differ substantially between CT26 or C26 tumor-bearing mice and healthy controls for the GAST (**Figure S2F**) or the TA (**Figure S2G**).

Despite the lack of a robust cachectic phenotype in mice bearing C26 and CT26 carcinomas at day 13 post-tumor cell inoculation, RT-qPCR analysis of the GAST muscle revealed that C26 mice already expressed more *Il6* compared to the CT26 group and also tended to have higher *Il6* mRNA than healthy controls (*p* = 0.119) (**Figure S3A**). Like late-stage cachexia, there was no change in muscle expression of other inflammatory cytokines including *Ccl2*, *Tnf*, and *Il1b* in C26 tumor-bearing mice (**Figure S3A**). Although not yet statistically different from healthy controls, C26 tumor-bearing mice did already display higher levels of immune cell markers including *Itgam* and *Mrc1* than the CT26 group (**Figure S3B**). Intriguingly, both tumor-bearing groups showed markedly reduced muscle expression of the atrophy related gene MuRF1 (*Trim63*) with a similar, yet not statistically significant response observed for the protein degradation marker Atrogin-1 (*Fbxo32*), indicating a potential compensatory strategy at this early cachexia stage (**Figure S3C**). Like our day 24 post-inoculation samples, muscle expression of *Ptgs2* was not altered in the muscle of CT26 or C26 tumor-bearing mice on day 13, but *Alox5* was already significantly increased in the C26 group (**Figure S3D**). In contrast, *Alox12* expression was found to be reduced relative to healthy controls in both CT26 and C26 tumor-bearing mice at this early stage, a response that was not observed at our later time-point (**Figure S3D**). Consistent with our day 24 data, we found that muscle 15-LOX protein was already significantly depleted in C26 tumor-bearing mice on day 13 of tumor growth, despite there not yet being an obvious change in muscle *Alox15* mRNA expression (**Figure S3E**). Complete uncropped western blots of 15-LOX are presented in **Figure S5**.

### C26 and CT26 Tumors Differently Modulate Skeletal Muscle Protein Turnover Pathways

To study the potential influence of C26 and CT26 tumors on regulating muscle protein turnover pathways, we determined the phosphorylation levels of key enzymes involved in protein synthesis signaling in GAST muscles via western blotting. Neither C26 nor CT26 tumor inoculation influenced the phosphorylation of protein kinase B (Akt) at either Thr308 or Ser473 (**Figure 6A**). C26 tumor-bearing mice had a trend towards increased phosphorylation of p70 ribosomal protein S6 kinase (p70S6K) at Thr389 (*p* = 0.079), but phosphorylation of p70S6K at Thr421/Ser424 rather tended to be decreased (*p* = 0.070) (**Figure 6B**). We also observed a modest yet statistically significant decrease in the phosphorylation of extracellular signal-regulated kinase 1/2 (ERK1/2, Thr202/Tyr204) in the C26 group relative to both healthy controls and CT26 mice (**Figure 6C**). Intriguingly, there was rather a significant increase in the phosphorylation of ribosomal protein S6 (S6) at both Ser235/236 and Ser240/244 sites in C26 tumor-bearing mice (**Figure 6D)**. In addition, we observed an increase in phosphorylation of the stress-activated protein kinase (SAPK) c-Jun N-terminal kinase (JNK) at Thr193/Tyr185 in CT26 but not C26 tumor-bearing mice (**Figure 6E)**. As opposed to the marked reduction seen at day 13 following tumor cell inoculation, there was rather a significant upregulation of *Fbxo32* in mice bearing C26 but not CT26 tumors (**Figure 6F**). Similarly, C26 tumors tended to induce an increase in *Trim63* (*p* = 0.090), while CT26 tumors did not influence muscle expression of *Trim63* (**Figure 6G**). Complete uncropped western blots of major phosphoproteins are presented in **Figure S6**.

**Figure 6.**
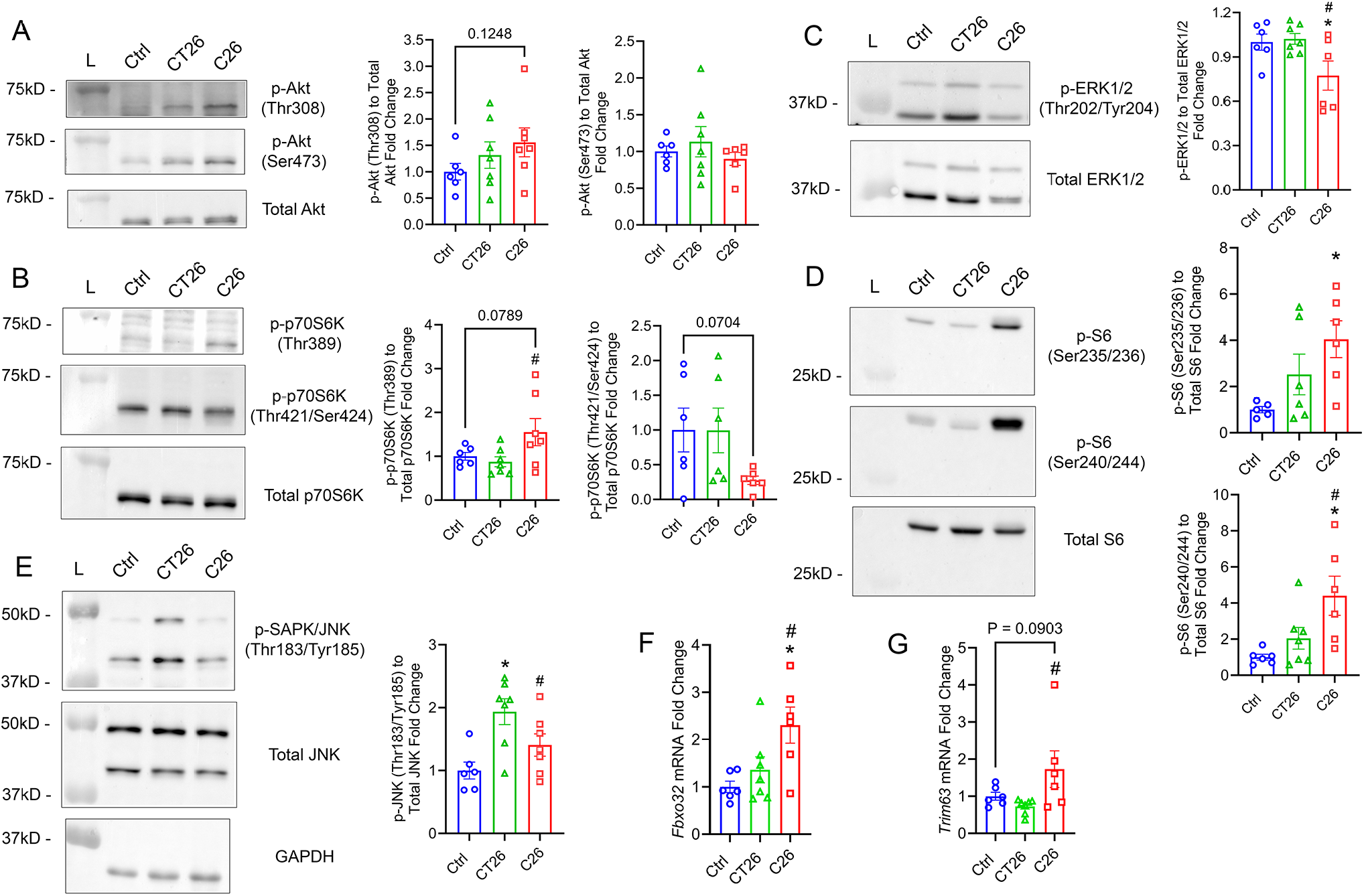
C26 and CT26 tumors exert different impacts on muscle protein synthesis and degradation pathways. Protein was extracted from GAST muscle homogenates, and western blot was used to determine the phosphorylation levels of **(A)** Akt at Thr308 and Ser473, **(B)** p70S6K at Thr389 and Thr421/Ser424, **(C)** ERK1/2 (Thr202/Tyr204), **(D)** S6 at Ser235/236 and Ser240/244, and **(E)** JNK (Thr183/Tyr185), with GAPDH used as a loading control. The mRNA expression level of the muscle-specific E3 ubiquitin ligases Atrogin-1 (*Fbxo32*) **(F)** and MuRF1 (*Trim63*) **(G)** were measured by RT-qPCR. Gene expression was normalized to the geometric mean of *Vcp*, *Emc7*, and *Gapdh*. Data are presented as the mean ± SEM, with dot plots showing raw data obtained from each individual mouse (biological replicates). Statistical significance was determined by one-way ANOVA followed by Fisher’s LSD post hoc tests. \**p* < 0.05 for difference compared to healthy controls, #*p* < 0.05 for difference between C26 and CT26 tumor-bearing mice.

### Differential Local Inflammatory Profile of C26 and CT26 Tumors

To further examine the inflammatory profile of C26 and CT26 tumors, we assessed the local gene expression of key cytokines that may potentially contribute to cachexia via their crosstalk with skeletal muscle. We found that C26 tumors obtained at 24 days post inoculation had significantly higher expression of *Il6* than CT26 tumors, indicating a hyper-inflammatory tumor profile (**Figure 7A**). However, C26 tumors expressed lower levels than CT26 tumors of certain other pro-inflammatory cytokines including *Ccl2*, *Tnf*, and *Lif* (**Figure 7A**). Moreover, there was no difference in the expression level of *Il1b* between CT26 and C26 tumors (**Figure 7A**). CT26 tumors exhibited a significantly higher level that C26 tumors of the pan myeloid cell marker *Itgam* as well as MΦ markers including *Cd68*, *Mrc1*, and *Arg1* (**Figure 7B**). C26 tumors also had a higher expression of *Ptgs2,* but a lower level of *Alox5,* when compared to CT26 tumors, while no difference was detected in the local expression of *Alox12* and *Alox15* (**Figure 7C**). Last but not least, we observed a significantly higher expression of the major stress hormone growth differentiating factor 15 (GDF15, *Gdf15)* in C26 compared to CT26 tumors, (**Figure 7D**).

**Figure 7.**
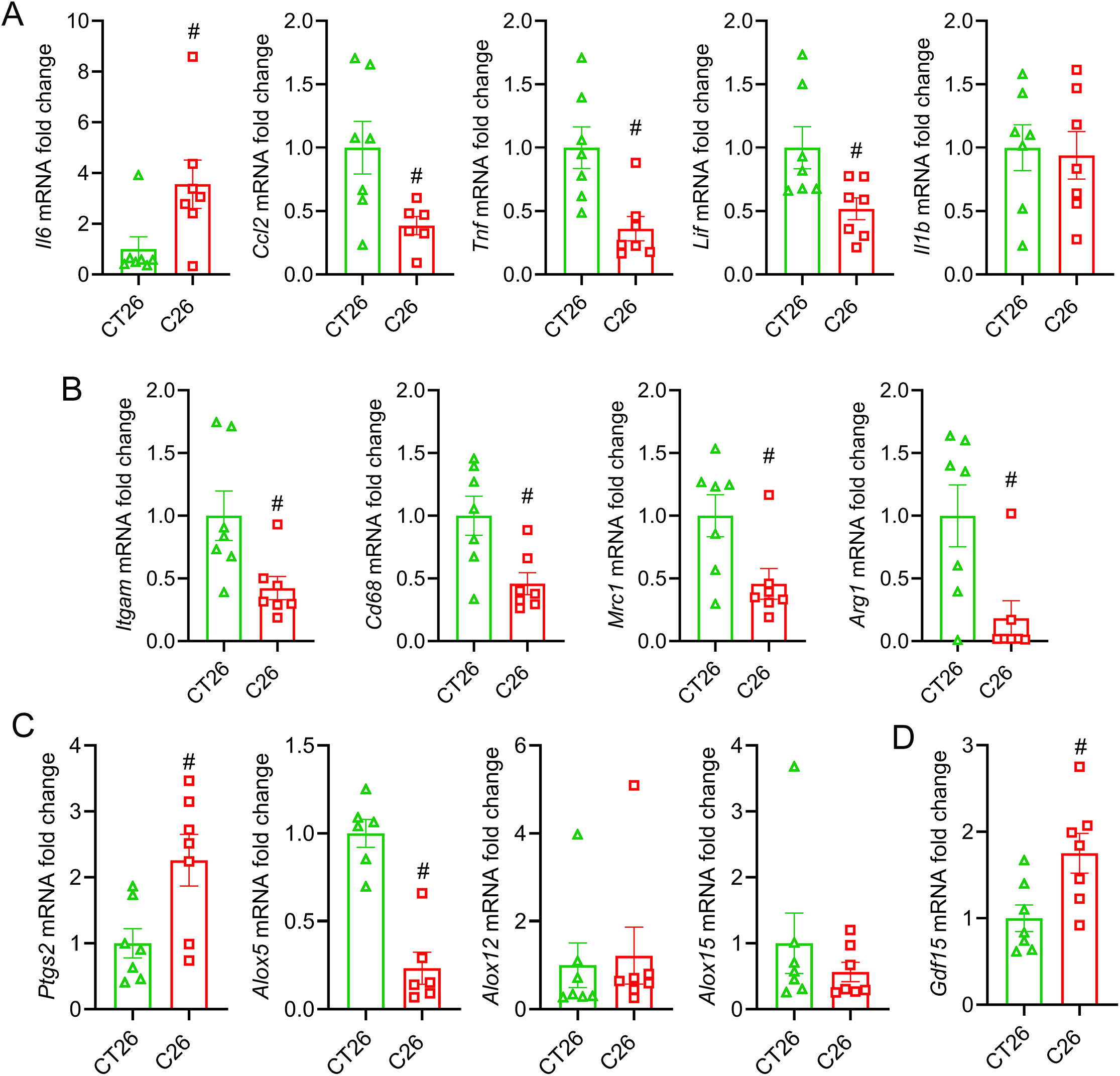
C26 and CT26 tumors have distinct inflammation-resolution gene profiles. RT-qPCR was used to determine the mRNA expression level of **(A)** pro-inflammatory cytokines (*Il6*, *Ccl2*, *Tnf*, *Lif*, and *Il1b*), **(B)** innate immune cell markers (*Itgam*, *Cd68*, *Mrc1*, and *Arg1***), (C)** lipid biosynthesis enzymes (*Ptgs2, Alox5*, *Alox12*, and *Alox15*), and **(D)** the stress hormone *Gdf15*. Gene expression was normalized to the geometric mean of *Vcp*, *Emc7*, and *Gapdh*. Data are presented as the mean ± SEM, with dot plots showing raw data obtained from each individual mouse (biological replicates). Statistical significance was determined by two-tailed unpaired t-tests. #*p* < 0.05 for difference between C26 and CT26 tumor-bearing mice.

A similar inflammatory profile was observed in tumor samples obtained in the earlier stages of cachexia. At 13 days post tumor cell inoculation, C26 tumors already tended to have higher local mRNA expression of *Il6* (*p* = 0.052) but lower levels of *Ccl2* (*p* = 0.068) and *Lif* (*p* = 0.153) when compared to CT26 tumors (**Figure S4A**). In addition, CT26 tumors already had higher mRNA expression than C26 tumors of innate immune cell markers including *Itgam*, *Cd68*, and *Mrc1*, with a similar pattern observed for *Arg1* (*p* = 0.120) (**Figure S4B**). Like our day 24 data set, C26 tumors also already tended to have a higher level of *Ptgs2* (*p* = 0.094) and a lower level of *Alox5* (*p* = 0.074) (**Figure S4C**). However, there was not yet any difference in the tumor-secreted level of *Gdf15* (**Figure S4D**).

## DISCUSSION

Colorectal cancer (CRC) cachexia involves involuntary loss of skeletal muscle mass and function, which could lower quality of life in cancer patients and reduce overall prognosis (1, 57). Although both the C26 and CT26 murine colon carcinoma cell lines have been used to model CRC cachexia, limited research has directly compared their relative potential to induce cancer cachexia. In the current study, we investigated the capacity of C26 and CT26 to induce CRC-associated cachexia in both *in vitro* and *in vivo* models. While both cell lines resulted in muscle atrophy *in vitro*, C26 tumors exhibited a higher capacity of inducing cachexia in a mouse model. We also show that C26 and CT26 could influence skeletal muscle dynamics through different biological mechanisms.

Previous research demonstrates that conditioned media from both C26 and CT26 carcinoma cells induces significant atrophy of murine C2C12 skeletal myotubes (58–61). A recent study used a co-culture model to investigate the crosstalk between C26 cells and C2C12 cells, showing that co-culture with C26 cells led to severe muscle atrophy (14). Using this muscle-cancer cell co-culture model, we tested the relative potency of C26 and CT26 cells to induce C2C12 atrophy at different seeding densities. We found that CT26 and C26 cells equivalently induced muscle atrophy in C2C12 myotubes, indicated by a significant decrease in myotube diameter. While CT26 cells led to a higher reduction in total myotube area, only C26 decreased the myogenin/DAPI ratio. As a crucial myogenic regulatory factor, myogenin drives the terminal differentiation of myoblasts and is critical for the expression of late muscle genes including muscle creatine kinase (62). It has been shown that dysfunctional myogenin interferes with myocyte fusion and leads to smaller fiber size *in vivo* (63). In addition to regulating muscle differentiation, myogenin also plays important roles in maintaining adult muscle fibers and regulating muscle stem cells (64). Our results indicate that while both C26 and CT26 cells induced muscle atrophy, C26 cells further impaired myogenin expression in mature C2C12 myotubes, which could contribute to additional defects in myotube fusion and maintenance.

On the other hand, C26 and CT26 cells exhibited differential capacity to induce cachexia in the mouse model. Although both CT26 and C26 tumor-bearing mice had a reduced tumor-free body weight, C26 tumors led to a higher loss of tumor-free lean mass and skeletal muscle mass. Moreover, while both tumor models exhibited a similar overall survival probability, the underlying clinical criteria necessitating humane euthanasia were profoundly distinct. While the primary driver for CT26 was the tumor burden, driver for C26 tumor-bearing mice was the severe cachexia (BCS < 2). Intriguingly, contrary to prior research, we show that both C26 and CT26 tumors led to reduced weight in GAST and TA, while a decrease in PLA, EDL, and SOL mass was only observed in C26 tumor-bearing mice (34). Differential changes of skeletal muscle mass are observed in different tumor models, and a higher reduction in muscle mass often indicates a more severe or late-stage cachexia (65, 66). We also observed muscle-specific changes in fiber morphology and composition of C26 tumor-bearing mice. The decrease in myofiber CSA of the predominantly slow-twitch SOL was driven mainly by an atrophy of type I fibers, while in the fast-twitch EDL there was a reduced type IIB fiber CSA while type I fiber size was rather simultaneously increased. In addition, we found that C26 tumors resulted in more severe muscle dysfunction. Consistent with previous research using animal models of cancer cachexia, C26 tumor-bearing mice in our study showed a deficit in force output coupled with a significant decrease in muscle mass (65, 67). Despite the observed decrease in muscle mass of TA and GAST, CT26 tumor-bearing mice did not exhibit loss of TA or GAST strength during force frequency testing. However, similar TA-specific force indicates a preserved intrinsic myofiber contractility in both groups of tumor-bearing mice, thus the reduced force output could be primarily driven by the more severe muscle atrophy in C26 tumor-bearing mice. When taken together, the reduced muscle mass yet preserved contractility in CT26 tumor-bearing mice could indicate milder symptoms of cancer cachexia compared to C26 tumors.

In addition to muscle mass, C26 tumor-bearing mice showed a higher reduction in fat mass compared to CT26 tumor-bearing mice. The decrease in adipose tissue in cancer cachexia could be attributed to increased lipolysis, white adipose tissue browning, and altered metabolism featured by higher energy expenditure (68–70). Previous research demonstrates a rewiring of adipose metabolism in C26-induced cachexia, upregulating the production of metabolites including sarcosine and thymidine (71). C26 tumor-bearing mice exhibited a higher ability to induce adipose wasting, which could contribute to the progression of skeletal muscle wasting. Future work could investigate the differential adipose metabolism in C26 and CT26 tumor-bearing mice and how it influences skeletal muscle growth in the two models.

To illustrate potential mechanisms by which C26 and CT26 differentially impact skeletal muscle mass and function, we examined the intramuscular innate immune responses in different muscle types. We observed a higher level of muscle inflammation in C26 tumor-bearing mice compared to CT26 in both the SOL and EDL muscle. Immunohistochemistry on SOL showed that C26 inoculation induced the infiltration of CD68^+^ and CD206^+^ MФ. Similarly, RT-qPCR also showed that C26 tumor-bearing mice had a significantly higher mRNA expression of the pan innate immune cell marker CD11b (*Itgam)* and the M2 macrophage marker CD206 (*Mrc1*) in the GAST. Previous studies have identified higher activity of tumor-associated macrophages (TAMs) in the tumor microenvironment, which promotes tumor proliferation and drives the production of cytokines such as IL-6 and CCL2 (72). It has also been shown that quadriceps from C26 tumor-bearing mice exhibited higher infiltration of both CXC chemokine receptor 1^+^ (CX3CR1^+^) MФ and anti-inflammatory polarized M2-like MФ (73). Although M2 MФ are often considered to be anti-inflammatory/pro-regenerative and play an essential role in the resolution of skeletal muscle inflammation, M2 MФ secreted factors may also exert pro-tumorigenic roles and promote muscle wasting in the context of cancer (74, 75). Similarly, M2 MФ have been shown to upregulate the TNF Receptor Associated Factor 6/NF-κB (TRAF6/p65) pathway in pancreatic cancer, which promotes the activation of Atrogin-1 and MuRF1 in skeletal muscle cells (76).

C26 tumor-bearing mice also exhibited a higher of intramuscular PMNs (Gr-1^+^ cells) in both SOL and EDL compared to CT26. Likewise, an elevation of both circulatory and intramuscular PMNs has been reported in C26 tumor-bearing mice, and such increase was not observed in CT26-bearing CD2F1 mice (34). Pathologically activated PMNs could potentially contribute to skeletal muscle wasting in cancer cachexia (77). Previously, research demonstrated that the polymorphonuclear myeloid-derived suppressor cells (PMN-MDSCs) promote skeletal muscle wasting via upregulation of the activin A signaling pathway in a mouse model of Lewis lung carcinoma (LLC) (78). Collectively, the differential recruitment of innate immune cells to skeletal muscle could contribute to the different cachectic effects of C26 and CT26 tumors.

Despite clear evidence of cellular inflammation of the SOL and EDL of C26 tumor bearing mice, we did not observe an obvious transcriptional change of classical pro-inflammatory cytokines or chemokines such as IL-6 (*Il6*), MCP-1 (*Ccl2*), TNFα (*Tnf*), and IL1β (*Il1b*) in the GAST muscle. Intriguingly, previous research has shown a differential expression of cytokines and chemokines in the TA and GAST muscle of C26 tumor-bearing mice. While the TA muscle showed a significant increase in the expression level of MCP-1 (*Ccl2*), C-X-C motif ligand 1 (CXCL1, *Cxcl1*), and CXCL2 (*Cxcl2*), the GAST only showed a massive increase in CXCL1 expression, revealing a differential inflammatory response to cancer progression in different muscle types (79). Importantly, RT-qPCR analysis of tumor tissues collected in the current study show that C26 tumors expressed a significantly higher local levels of IL-6 (*Il6*), one of the primary drivers of CRC cachexia, which is thought to contribute to the elevated immune infiltration of skeletal muscle. Indeed, it has been previously established that cachectic factors such as IL-6 are primarily secreted by tumor rather than muscle cells (80). The divergent regulation of IL-6 secretion in tumor tissues provides a potential mechanism responsible for the more severe cachectic symptoms in C26 tumor-bearing mice. Interestingly, we found that C26 tumors generally had lower expression of other pro-inflammatory cytokines responsible for immune cell recruitment, such as MCP-1 (*Ccl2*) and TNFα (*Tnf*). Similarly, C26 tumors expressed lower levels of innate immune cell markers including CD11b (*Itgam*), CD68 (*Cd68*), and arginase 1 (*Arg1*), indicating an immune-desert phenotype of C26 tumors. This finding is consistent with the previous research showing that CT26 tumors contain more infiltrating immune cells while C26 tumors display relatively greater immune evasion (29). Moreover, we observed a higher expression of COX-2 (*Ptgs2*) and GDF15 (*Gdf15*) in C26 tumor tissues compared to CT26, indicating a COX-dominant pro-tumorigenic tumor microenvironment and an elevated cachexia signaling in C26 tumors. The elevated expression of *Gdf15* could also contribute to the lower food intake and higher adipose wasting observed in C26 tumor-bearing mice (81).

We found that C26 and CT26 tumors also differentially mediated the expression of lipid biosynthesis enzymes in cachectic skeletal muscle. C26 tumors significantly increased muscle mRNA expression of 5-LOX (*Alox5*) while no such change was found in CT26 tumor-bearing mice. 5-LOX is the essential enzyme involved in the production of eicosanoids derived from the omega-6 arachidonic acid (ARA, 20:4n-6), including the leukotrienes (82). In an acute model of myotoxin-induced skeletal muscle injury, it has been suggested that excessive 5-LOX activity may result in increased inflammation and impaired myofiber regeneration (83). Elevated 5-LOX activity has also been reported to drive myotube atrophy induced by glucocorticoids through promoting local production of the pro-inflammatory eicosanoid leukotriene B_4_ (LTB_4_) and activating the nuclear factor kappa B (NF-κB) pathway (84). Additionally, it has been shown that the 4-weeks of treatment with the pharmacological 5-LOX inhibitor malotilate could alleviate skeletal muscle wasting in aging mice (84). On the other hand, we recently found that both malotilate and the 5-LOX activating protein (FLAP) inhibitor MK886 markedly interfered with the myogenic differentiation of C2C12 myoblasts (55). Therefore, the 5-LOX pathway may play a dual role in skeletal muscle plasticity under different circumstances. Nevertheless, our data suggests that chronically increased 5-LOX activity may contribute to skeletal muscle wasting in C26 tumor bearing mice.

Unlike 5-LOX, we observed a tumor-specific suppression of leukocyte-type 12/15-LOX (*Alox15*) in CT26 and C26 tumor-bearing mice, which is a key enzyme mediating SPM production and the resolution of inflammation (85). A reduced expression of 15-LOX-1, the human ortholog of murine 12/15-LOX, has been reported in human colorectal and pancreatic adenocarcinoma cell lines, which contributes to tumor survival and dysregulated tumor growth (86–88). However, studies on the influence of cancer cachexia on the expression of skeletal muscle 12/15-LOX have been limited. Here, we show that at late stage of cachexia, C26 tumors markedly diminished both the gene and protein level of 12/15-LOX in GAST. On the other hand, while CT26 tumors also reduced the mRNA expression of 12/15-LOX, no protein deficiency was found in CT26 muscles, indicating potential post-transcriptional mechanisms to compensate for 12/15-LOX function in CT26 tumor-bearing mice. Similar findings of diminished muscle 15-LOX protein expression were also observed in the early stages of cachexia at day 13 following tumor inoculation prior to the onset of cancer-associated loss of muscle mass and strength. This suggests that deregulation of skeletal muscle inflammation-resolution circuits is an important underlying mechanism that precedes the onset of muscle wasting, rather than simply being a downstream consequence of advanced stage cachexia.

It was recently reported that genetic deficiency of 12/15-LOX blocked the class switch of bioactive lipid mediators following acute muscle injury, resulting in impaired skeletal muscle regeneration following acute injury (54, 55). Transgenic mice lacking 12/15-LOX enzyme (*Alox15*^-/-^) also exhibited chronic basal skeletal muscle inflammation featured by increased infiltration of PMNs and MФ (55). The diminished expression of 12/15-LOX in C26 tumor-bearing mice at both early and late cachexia stages could contribute to the progression of muscle wasting through mediating inflammatory responses. Collectively, the distinct regulation of 5-LOX and 12/15-LOX expression in skeletal muscles could potentially contribute to the differential intramuscular immune responses and cachectic effects in CT26 and C26 tumor-bearing mice. Intriguingly, in LLC-induced cancer cachexia, despite a lower expression of 12/15-LOX protein in the GAST muscle, the production of oxylipins derived from 12/15-LOX was apparently elevated, which has been suggested to contribute to progression of a cachexia phenotype by promoting oxidative stress (89). This could suggest a distinct pathophysiology of cancer cachexia induced by C26 and LLC. Further research should assess the intramuscular concentrations and biological roles of various bioactive lipid mediators derived from the 5-LOX and 12/15-LOX pathways in C26 and CT26 mice.

We also show that C26 and CT26 tumors differentially regulated signaling pathways involved in protein synthesis, particularly the Akt/mTOR/p70S6K pathway, which plays a crucial role in regulating muscle protein synthesis (90, 91). Although no significant change was observed in the phosphorylation level of Akt, C26 tumor-bearing mice tended to have an increased phosphorylation level of p70S6K at Thr389, whereas CT26 tumor-bearing mice did not. Meanwhile, we also found that phosphorylation level of ERK1/2 was repressed in C26 tumor-bearing mice, which phosphorylates the autoinhibitory site Thr421/Ser424 of p70S6K (92). Indeed, we observed a concurrent downregulation of phosphorylation of p70S6K at Thr421/Ser424. Intriguingly, we observed a simultaneous elevation of phosphorylation of ribosomal protein S6 at both Ser235/236 and Ser240/244, a well-established downstream effector of the mTOR/p70S6K pathway, in C26 tumor-bearing mice despite a significant decrease in muscle mass. These activities together map a hyperactive mTORC1 signaling bypassing ERK1/2 activation to activate p70S6K and downstream effectors, potentially in response to the release of amino acids following higher protein degradation (93–96).

Paradoxically, some studies have shown that p70S6K/S6 signaling is largely impaired in colorectal cancer, indicated by the decreased phosphorylation level of p70S6K, S6, or 4E-BP1, which leads to decreased protein synthesis (23, 97, 98). However, a recent study showed that C26 tumor inoculation did not necessarily decrease the phosphorylation level of p70S6K or S6 despite significant loss of skeletal muscle mass (99). Other research has also observed S6 hyperactivation combined with severe muscle atrophy under pathological stress, showing the decoupling of S6 phosphorylation and actual muscle metabolic outcome (100, 101). Meanwhile, the hyperactivation of S6 potentially contributes to the lack of changes observed in Akt phosphorylation via the negative feedback regulation of the upstream signal protein (102).

Notably, CT26 but not C26 tumor-bearing mice had a significantly upregulated phosphorylation of JNK at Thr183/Tyr185, which is another subfamily of MAPK (103). The JNK/c-Jun pathway could be associated with increased potential of metastasis and cancer stemness (104, 105). In the context of pancreatic cancer cachexia, it has been suggested that the activation of JNK signaling contributed to muscle wasting by activating the muscle-specific E3 ubiquitin ligases Atrogin-1 (*Fbxo32*) and MuRF1 (*Trim63*) (106). However, our model shows that despite the increased JNK phosphorylation in both CT26 and C26 mice, the mRNA expression of these muscle-specific E3 ubiquitin ligases was increased only in mice bearing C26 tumors. This result suggests other regulatory mechanisms of protein degradation pathway in CT26 tumor-bearing mice that counteracts the activation of JNK signaling. Furthermore, the greater mRNA expression of muscle-specific E3 ubiquitin ligases in C26 vs CT26 mice indicates a potentially higher protein degradation induced by C26 tumors via a JNK independent mechanism.

One limitation of this study is that circulating levels of pro-inflammatory cytokines were not measured. Despite no change in cytokine expression was detected in GAST muscles, tumor-secreted cytokines released to the plasma could contribute to systemic inflammation and cancer cachexia. Although we assessed and compared local expression of tumor-secreted cytokines as a proxy measure, we could not compare the level back to healthy controls. We also did not examine the systemic or intramuscular profile of pro-inflammatory and anti-inflammatory/pro-resolving lipid mediators derived from various LOX-dependent pathways. Future studies should assess whether diminished 15-LOX and upregulated 5-LOX expression in skeletal muscle tissue of C26 tumor-bearing mice impact downstream concentrations of respective bioactive lipid mediators. Moreover, while recent studies indicate that male mice may be more prone to C26-induced cachexia, we used female BALB/c mice to compare the intrinsic difference of C26 and CT26 carcinoma (107, 108). Hence, future studies should investigate whether the differential effects of C26 and CT26 cells are impacted by sex. Further investigation could also assess the potential time-dependent changes in cytokine production in skeletal muscles. In addition, while we assessed the phosphorylation activity of major enzymes involved in protein synthesis and measured the expression of muscle-specific E3 ubiquitin ligases, the rate of protein synthesis or proteolysis was not directly measured. While our primary focus in the current study was the dynamics of skeletal muscle wasting, it could be worth examining the local cellular and molecular differences between C26 and CT26 tumors as well as how they differentially regulate adipose tissue wasting. Finally, future research could examine how muscle metabolism is mediated in C26 and CT26 tumor-bearing mice. Nevertheless, this study addressed a critical gap in preclinical research on cancer cachexia by illustrating the differential cachectic potential of CT26 and C26 cell lines *in vitro* and *in vivo*, which provides evidence for choosing appropriate cancer cachexia models in future studies. Our findings also indicate that the clinical severity of cachexia could be heavily determined by the differential crosstalk between tumor, muscle, and the host immune system, and future translational studies should consider the heterogeneity of patients with cancer cachexia to identify the potential difference in clinical molecular profile.

## CONCLUSION

In the current study, we found that both C26 and CT26 colon carcinoma cells induced skeletal muscle cell atrophy in a muscle-cancer co-culture model. However, in a mouse model of tumor inoculation, while CT26 tumors resulted in decreases in tumor-free body weight and muscle mass in GAST and TA, C26 inoculation caused a more significant loss of lean mass and a more severe impairment of skeletal muscle function. We show that C26 tumors also led to a higher level of intramuscular inflammation than CT26, and C26 muscles had an increased expression of 5-LOX and a repressed expression of 15-LOX. Moreover, the two cell lines differentially regulate protein turnover pathways. Despite the upregulated activity of JNK signaling, CT26 exhibited a milder cachexia phenotype. The differential pathophysiology of C26 and CT26 cancer cachexia highlights the importance of evaluating both tumor-secreted factors and intramuscular immune profile during therapeutic interventions.

## ACKNOWLEDGEMENTS

The MF20 (developed by Fischman, D.A.) and F5D (developed by Wright, W.E.) monoclonal antibodies were obtained from the Developmental Studies Hybridoma Bank (DSHB), created by the NICHD of the NIH and maintained at The University of Iowa, Department of Biology, Iowa City, IA 52242. We thank Dr. Keehong Kim (Purdue University) for the generous gift of the C26 cell line.

## AUTHOR CONTRIBUTIONS

**Xinyue Lu:** Conceptualization, Methodology, Validation, Formal analysis, Investigation, Writing – original draft, Writing – review & editing, Visualization. **Hanan Tlais:** Investigation, Formal analysis, Writing – review & editing. **Hamood Rehman:** Methodology, Software, Investigation, Writing – review & editing. **Ashtyn N. Martens:** Investigation, Formal analysis, Writing – review & editing. **Abigail L. Hartz:** Investigation, Formal analysis, Writing – review & editing. **Charles P. Olson**: Investigation, Formal analysis, Writing – review & editing. **Nina R. O’Tain**: Investigation, Formal analysis, Writing – review & editing. **Vandré C. Figueiredo:** Methodology, Formal analysis, Writing – review & editing, Visualization, Supervision. **Tingting Ju:** Methodology, Formal analysis, Resources, Writing – review & editing, Supervision, Funding acquisition. **James F. Markworth:** Conceptualization, Methodology, Software, Formal analysis, Resources, Writing – review & editing, Visualization, Supervision, Project administration, Funding acquisition.

---

**Figure S1.**
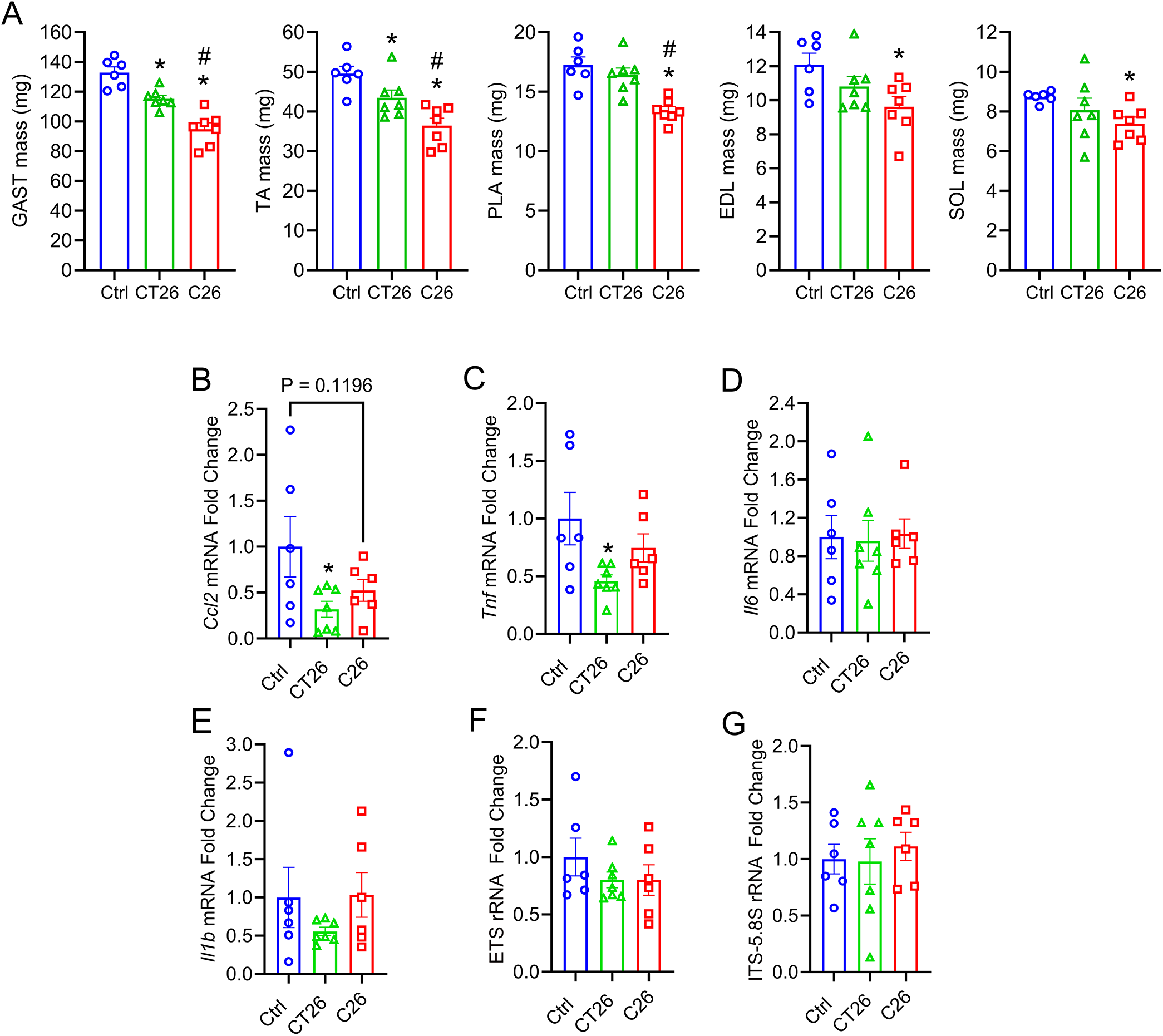
Terminal muscle mass and mRNA expression of pro-inflammatory cytokines and enzymes mediating ribosome biogenesis in skeletal muscle. (**A**) Final absolute mass of major hindlimb muscles including the gastrocnemius (GAST), tibialis anterior (TA), extensor digitorum longus (EDL), plantaris (PLA), and soleus (SOL). (**B-G**) RT-qPCR was used to determine the mRNA expression level of *Ccl2* (B), *Tnf* (C), *Il6* (D), and *Il1b* (E) in GAST muscles. The rRNA fold change of ETS **(F)** and ITS-5.8S **(G)** was also measured in GAST muscles of healthy controls, CT26 tumor-bearing mice, and C26 tumor-bearing mice. Gene expression was normalized to the geometric mean of *Vcp*, *Emc7*, and *Gapdh*. Data are presented as the mean ± SEM, with dot plots showing raw data obtained from each individual mouse (biological replicates). Statistical significance was determined by one-way ANOVA followed by Fisher’s LSD post hoc tests. \**p* < 0.05 for difference compared to healthy controls, #*p* < 0.05 for difference between C26 and CT26 tumor-bearing mice.

**Figure S2.**
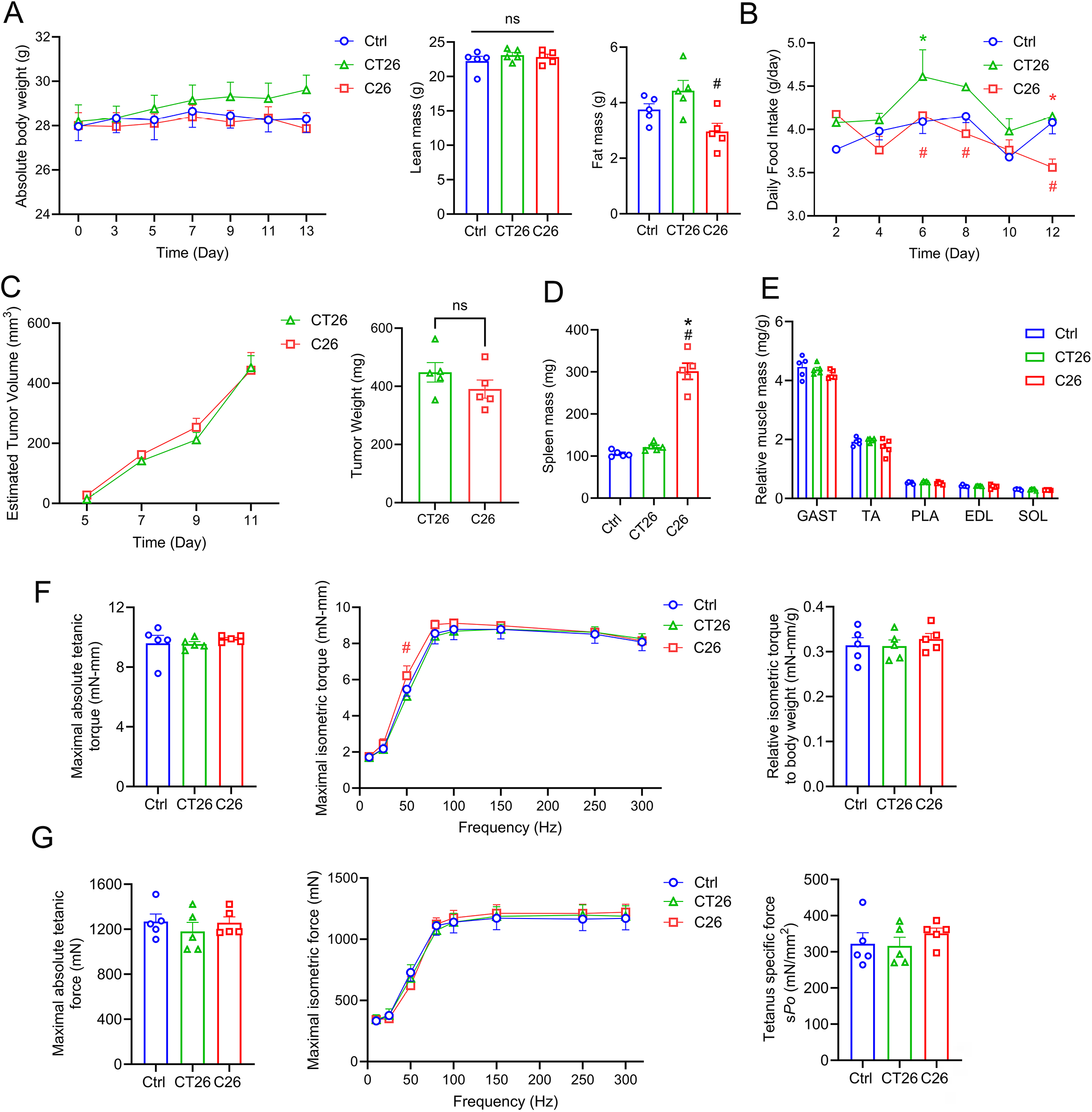
Absence of cachectic phenotypes in C26 or CT26 tumor-bearing mice at early cancer stage. Female BALB/c mice were inoculated with saline (*n* = 5), 5 × 10^4^ CT26 cells (*n* = 5), or 5 × 10^4^ C26 cells (*n* = 5). **(A)** Body weight was recorded every two days up to day 13 post inoculation, and absolute lean and fat mass was measured by EchoMRI one day prior to euthanasia. **(B)** Average food intake was recorded every two days up to day 12. **(C)** Tumor volume was estimated from tumor diameters and final tumor mass was weighed after dissection on the day of euthanasia. **(D)** Spleen weight of healthy, CT26 and C26 tumor-bearing mice was weighed after dissection. **(E)** Muscle mass of GAST, TA, PLA, EDL, and SOL expressed relative to initial body weight. **(F)** Maximal tetanic torque of plantar flexors at 150 Hz, isometric torque at increasing frequencies from 10-300 Hz, and relative tetanic torque when normalized to terminal body weight. **(G)** Absolute maximal tetanic force of TA at 150 Hz, isometric force at increasing frequencies from 10-300 Hz, and TA-specific maximal tetanic force (*sP_o_*). Data are presented as the mean ± SEM, with dot plots showing raw data obtained from each individual mouse (biological replicates). Statistical significance was determined by one-way ANOVA (bar charts) or two-way ANOVA (line graphs) followed by Fisher’s LSD post hoc tests. \**p* < 0.05 for difference compared to healthy controls, #*p* < 0.05 for difference between C26 and CT26 tumor-bearing mice.

**Figure S3.**
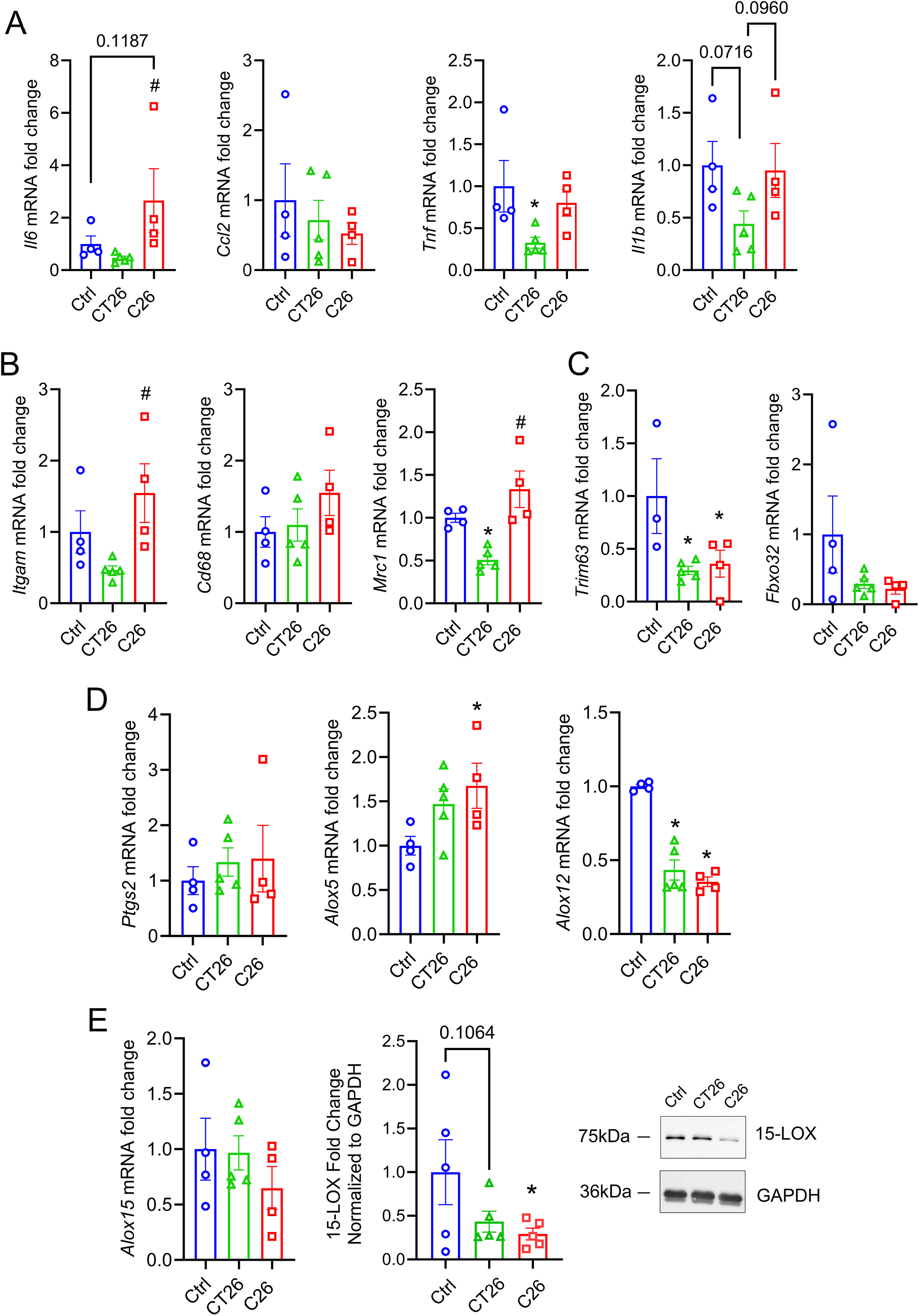
Skeletal muscle inflammation-resolution gene profile at an early cancer cachexia stage. The mRNA expression levels of **(A)** pro-inflammatory cytokines (*Il6*, *Ccl2*, *Tnf*, *Lif*, and *Il1b*), **(B)** immune cell markers (*Itgam*, *Cd68*, and *Mrc1*), **(C)** muscle-specific E3 ubiquitin ligases (*Trim63* and *Fbxo32*) and **(D)** lipid biosynthesis enzymes including *Ptgs2, Alox5*, and *Alox12* were determined in the GAST muscle by RT-qPCR. **(E)** mRNA and protein expression of 15-LOX were assessed by RT-qPCR and western blot, respectively. Gene expression was normalized to the geometric mean of *Vcp*, *Emc7*, and *Gapdh*. Data are presented as the mean ± SEM, with dot plots showing raw data obtained from each individual mouse (biological replicates). Statistical significance was determined by one-way ANOVA followed by Fisher’s LSD post hoc tests. \**p* < 0.05 for difference compared to healthy controls, #*p* < 0.05 for difference between C26 and CT26 tumor-bearing mice.

**Figure S4.**
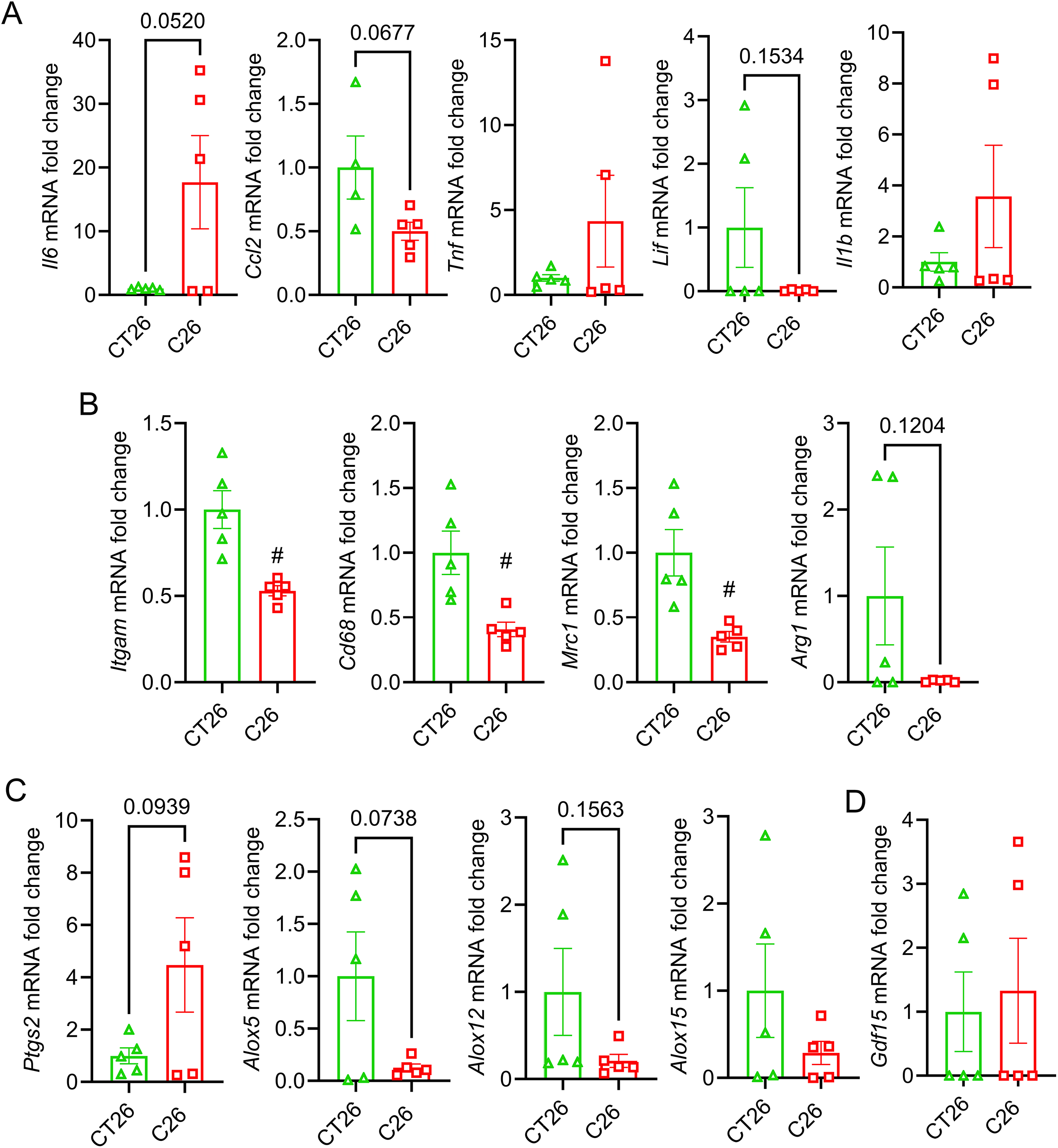
Tumor gene profile on day 13 post-inoculation of C26 and CT26 tumors. RT-qPCR was used to assess the mRNA expression level of target genes in both C26 and CT26 tumors on day 13 following injection of carcinoma cells, including **(A)** the pro-inflammatory cytokines (*Il6*, *Ccl2*, *Tnf*, *Lif*, and *Il1b*), **(B)** immune cell markers (*Itgam*, *Cd68*, *Mrc1*, and *Arg1***), (C)** lipid biosynthesis enzymes (*Ptgs2, Alox5*, *Alox12*, and *Alox15*), and **(D)** stress hormone *Gdf15*. Gene expression was normalized to the geometric mean of *Vcp*, *Emc7*, and *Gapdh*. Data are presented as the mean ± SEM, with dot plots showing raw data obtained from each individual mouse (biological replicates). Statistical significance was determined by two-tailed unpaired t-tests. #*p* < 0.05 for difference between C26 and CT26 tumor-bearing mice.

**Figure S5.**
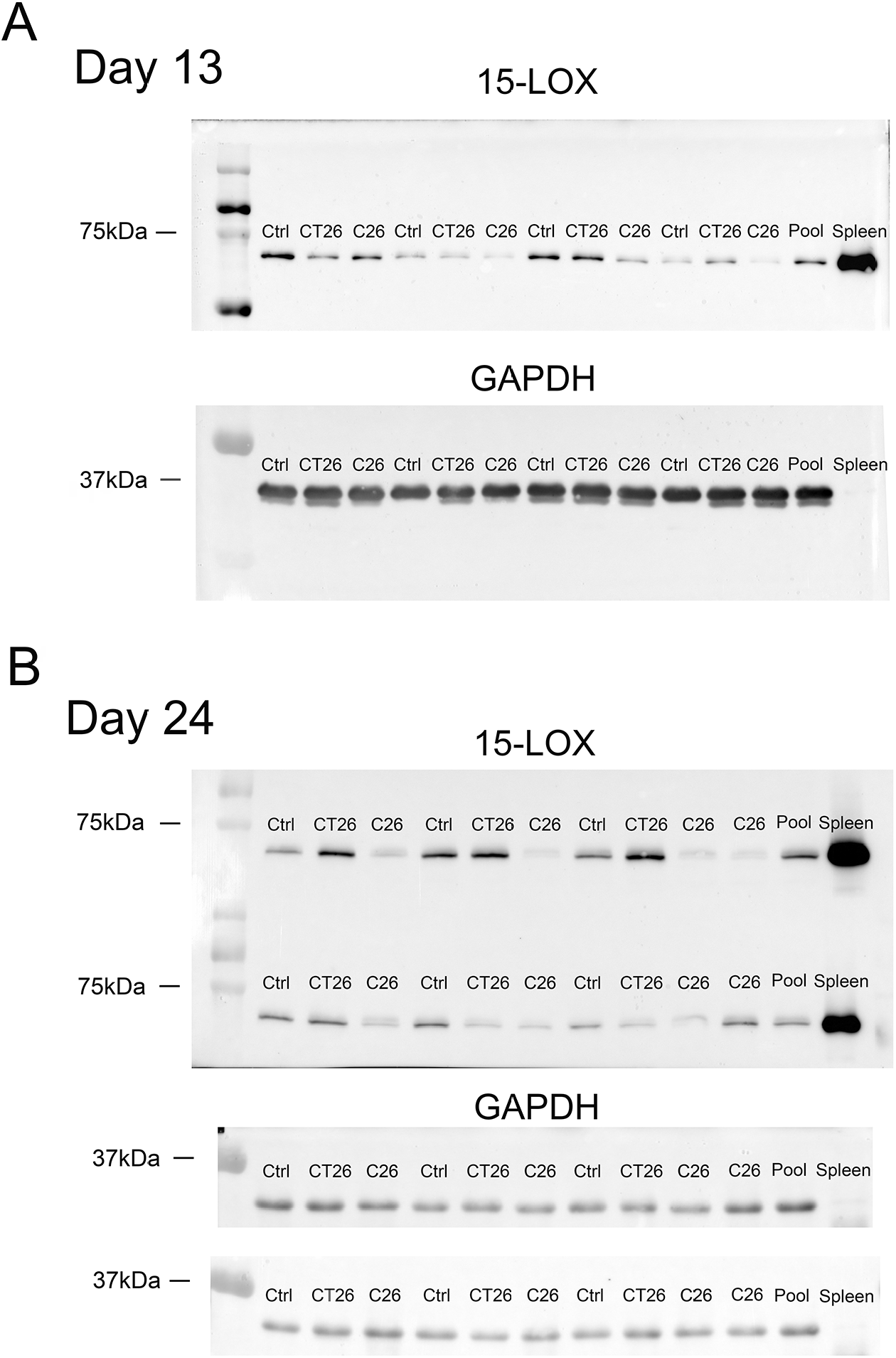
Western blots analysis of 15-LOX expression. Full western blots of GAST muscle protein homogenate from all the samples in Ctrl, CT26, and C26 groups obtained at both early (day 13) and late (day 24) cachexia stages. The protein expression of 15-LOX at day 13 **(A)** and day 24 **(B)** is shown together with GAPDH bands as a loading control.

**Figure S6.**
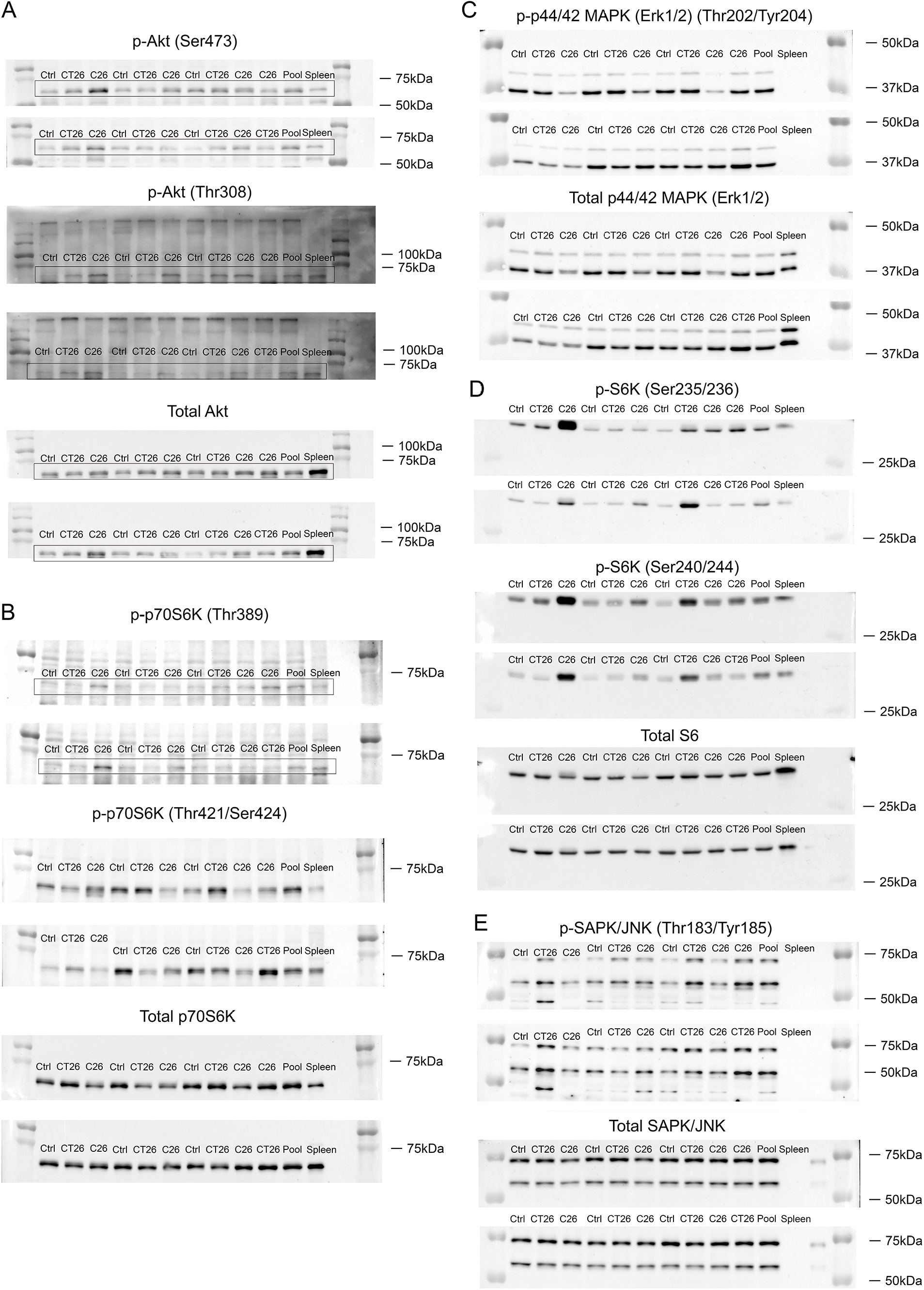
Western blots analysis of protein synthesis signaling. Full western blots of GAST muscle protein homogenate from all the samples obtained at the late cachexia stage (day 24) in the Ctrl (*n* = 6), CT26 (*n* = 7), and C26 (*n* = 7) groups. Target proteins include **(A)** p-Akt (Ser473), p-Akt (Thr308), and total Akt, **(B)** p-p70S6K (Thr389), p-p70S6K (Thr421/Ser414), and total p70S6K, **(C)** p-ERK1/2 (Thr202/Tyr204) and total ERK1/2, **(D)**, p-S6 (Ser235/236), p-S6 (Ser240/244), and total S6, and **(E)** p-JNK (Thr183/Tyr185) and total JNK.

